# Target Capture Sequencing Unravels *Rubus* Evolution

**DOI:** 10.1101/703926

**Authors:** Katherine A. Carter, Aaron Liston, Nahla V. Bassil, Lawrence A. Alice, Jill M. Bushakra, Brittany L. Sutherland, Todd C. Mockler, Douglas W. Bryant, Kim E. Hummer

## Abstract

**Background:** *Rubus* (Rosaceae) comprises more than 500 species with additional commercially cultivated raspberries and blackberries. The most recent (> 100 years old) global taxonomic treatment of the genus defined 12 subgenera; two subgenera were subsequently described and some species were rearranged. Intra- and interspecific ploidy levels and hybridization make phylogenetic estimation of *Rubus* challenging. Our objectives were to: estimate the phylogeny of 94 geographically diverse species and 3 cultivars using chloroplast DNA sequences and target capture of approximately 1,000 low copy nuclear genes; estimate divergence times between major *Rubus* clades; and examine the historical biogeography of species diversification.

**Results:** Target capture sequencing identified eight major groups within *Rubus*. Subgenus *Orobatus* and Subg. *Anoplobatus* were monophyletic, while other recognized subgenera were para- or polyphyletic. Multiple hybridization events likely occurred across the phylogeny at subgeneric levels, *e*.*g*., Subg. *Rubus* (blackberries) × Subg. *Idaeobatus* (raspberries) and Subg. *Idaeobatus* × Subg. *Cylactis* (Arctic berries) hybrids. The raspberry heritage within known cultivated blackberry hybrids was confirmed. The most recent common ancestor of the genus was most likely distributed in North America. Multiple distribution events occurred during the Miocene (about 20 Ma) from North America into Asia and Europe across the Bering land bridge and southward crossing the Panamanian Isthmus. *Rubus* species diversified greatly in Asia during the Miocene.

**Conclusions:** *Rubus* taxonomy does not reflect phylogenetic relationships and subgeneric revision is warranted. Target capture sequencing confirmed that most subgenera are para- or polyphyletic. The most recent common ancestor migrated from North America towards Asia, Europe, and Central and South America early in the Miocene then diversified. Ancestors of the genus *Rubus* may have migrated to Oceania by long distance bird dispersal. This phylogeny presents a roadmap for further *Rubus* taxonomic and phylogenetic research.

## Background

The plant genus *Rubus* (Rosaceae), contains a conservative estimate of more than 500 species (1) and thousands of cultivars. The annual production of the cultivated brambles (raspberries and blackberries), is economically significant for more than 43 countries (2). Crop wild relatives of this genus contribute to broadening the gene pools for breeding programs to improve these nutritious berry crops.

Varying intra- and interspecific ploidy levels (diploid, 2*n* = 2*x* = 14 to dodecaploid, 2*n* = 12*x* = 84, plus aneuploids), and hybridization (3–8) make phylogenetic estimation challenging. Focke’s worldwide taxonomic treatment of *Rubus* (9–11), defined 12 subgenera (Table 1). Subg. *Rubus* (= *Eubatus* Focke), *Idaeobatus*, and *Malachobatus* contain the most species with > 300 species/microspecies for subg. *Rubus*, 88 species for subg. *Idaeobatus* and 92 species for subg. *Malachobatus* (3, 12).

**Table 1.** Accessions of *Rubus* species and outgroup (*Waldsteinia fragarioides*) used in this study. Species marked with an asterisk in the “Species” column did not sequence well and were not included in the results. Subgenera classifications in Focke and the USDA GRIN network are reported. Subgenera marked with an asterisk in the “USDA GRIN Subgenus Classification” column are not listed in GRIN. Focke subg. *Eubatus* has been renamed to subg. *Rubus*. Current classifications were curated from other publications (13, 92–94). Herbarium vouchers with collector, number, and herbarium (63) or PI numbers for accessions of plants housed in the living collection at USDA NCGR Corvallis are given. MOR refers to the living collection at Morton Arboretum, Lisle, IL. HPDL refers to the Native Hawaiian Plants DNA library (65). The geographic origin for each accession is listed by continent or region. Ploidy data was collected from flow cytometry data, multiple publications, and the Missouri Botanical Garden index of plant chromosome number database (4, 5, 95, 96). Eight major phylogenetic groups were identified in nuclear sequence analyses. The group in which each species is found is listed.

| Species | Ploidy | USDA GRIN<br>Subgenus<br>Classification | Focke Subgenus<br>Classification | Region of<br>Origin | Group (1-8) | Voucher |
| --- | --- | --- | --- | --- | --- | --- |
| *R. deliciosus* Torr. | 2x | *Anoplobatus* | *Anoplobatus* | North<br>America | 2 | PI 553184/CRUB<br>1021.001 |
| *R. odoratus* L. | 2x | *Anoplobatus* | *Anoplobatus* | North America | 2 | Alice R14, MAINE |
| *R. parviflorus* Nutt. | 2x | *Anoplobatus* | *Anoplobatus* | North America | 2 | PI 553785/CRUB 13.001 |
| *R. trilobus* Thunb. | 2x | *Anoplobatus* | *Anoplobatus* | South America | 2 | Ruiz 889, MO |
| *R. calycinus* Wall. Ex D. Don | 6x | *Chamaebatus* | *Chamaebatus* | Asia | 5 | Alice, Sutherland and Dorji from Bhutan in Alice, Sutherland and Dorji (2008) from Rubus-Ribes 2005. Now Vouchered at WKU 04-07 |
| *R. nivalis* Douglas | 2x | *Chamaebatus* | *Chamaebatus* | North America | 8 | PI 679726/CRUB 1374.001 PL |
| *R. pectinellus* Maxim.* | 6x | *Chamaebatus* | *Chamaebatus* | Asia | n/a | Jutila & Fujino 680, MO |
| *R. pectinarioides* | 4x | *Chamaebatus* * | n/a | Asia | 5 | Alice, Sutherland and Dorji from Bhutan in Alice, Sutherland and Dorji (2008) from Rubus-Ribes 2005. Vouchered WKU 04-25 |
| *R. sengorensis* | 4x | *Chamaebatus* * | n/a | Asia | 5 | Alice, Sutherland and Dorji from Bhutan in Alice, Sutherland and Dorji (2008) from Rubus-Ribes 2005. |
|  |  |  |  |  |  | Vouchered WKU 04-33 |
| *R. chamaemorus* L. | 8x | *Chamaemorus* | *Chamaemorus* | North America/Northern Europe | 1 | Alice R17, MAINE |
| *R. geoides* Sm. | 4x | *Comaropsis* | *Comaropsis* | South America | 8 | Dudley et al. 1538a, MO |
| *R. arcticus* L. | 2x | *Cylactis* | *Cylactis* | North America/Northern Europe | 3 | T. Eriksson 701, S |
| *R. humulifolius* C. A. Mey. | 4x | *Cylactis* | *Cylactis* | Asia | 4 | PI 553242/CRUB 1173.001 PL |
| *R. saxatilis* L. | 4x | *Cylactis* | *Cylactis* | Europe/Asia | 7 | PI 370230/CRUB 918.001 PL |
| *R. lasiococcus* A. Gray | 2x | *Cylactis* | *Dalibarda* | North America | 1 | Merello et al. 827, MO |
| *R. pedatus* Sm. | 2x | *Cylactis* | *Dalibarda* | North America/Asia | 1 | Alice 96-1, MAINE |
| *R. fockeanus* Kurz | 4x | *Cylactis* | *Dalibarda* | Asia | 5 | PI 606537/CRUB 1960.000 SD |
| *R. pubescens* Raf. | 2x | *Cylactis* | *Eubatus* | North America | 3 | Alice R15, MAINE |
| *R. treutleri* Hook. f. | 4x | *Dalibardastrum* | *Dalibardastrum* | Asia | 5 | Alice, Sutherland and Dorji from Bhutan in Alice, Sutherland and Dorji (2008) from Rubus-Ribes 2005. |
|  |  |  |  |  |  | Now Vouchered at<br>WKU 04-09 |
| *R. tricolor* Focke | 4x | *Dalibardastrum* | *Dalibardastrum* | Asia | 5 | Alice 97-2, MAINE |
| *R. amphidasys* Focke | 6x | *Dalibardastrum* | *Malachobatus* | Asia | 5 | PI 618397/CRUB<br>1693.001 PL |
| *R. nepalensis* (Hook.f)<br>Kuntze | 4x | *Dalibardastrum* | n/a | Asia | 5 | Alice 97-1, MAINE |
| *R. gunnianus* Hook. | 4x | *Diemenicus* | *Dalibarda* | Australia | 8 | Wells 96-1, MAINE |
| *R. trifidus* Thunb. | 2x | *Idaeobatus* | *Anoplobatus* | Asia | 4 | PI 554051/CRUB 3.001<br>PL |
| *R. parvifolius* L. | 2x | *Idaeobatus* | *Eubatus* | Asia | 7 | PI 553813/CRUB 5.001<br>PL |
| *R. hawaiiensis* A. Gray | 2x | *Idaeobatus* | *Idaeobatus* | North<br>America<br>(Hawaii) | 3 | PI 553214/CRUB<br>399.001 PL |
| *R. spectabilis* Pursh | 2x | *Idaeobatus* | *Idaeobatus* | North<br>America | 3 | PI 553980/CRUB 4.001<br>PL |
| *R. crataegifolius*<br>Bunge | 2x | *Idaeobatus* | *Idaeobatus* | Asia | 4 | PI 553173/CRUB<br>16.001 PL |
| *R. ellipticus* Sm. | 2x | *Idaeobatus* | *Idaeobatus* | Asia | 4 | PI 553190/CRUB<br>1052.001 PL |
| *R. illecebrosus* Focke | 2x | *Idaeobatus* | *Idaeobatus* | Asia | 4 | PI 553643/CRUB<br>838.001 PL |
| *R. palmatus* Thunb. | 2x | *Idaeobatus* | *Idaeobatus* | Asia | 4 | PI 553782/CRUB 2.002<br>PL |
| *R. rosifolius* Sm. | 2x | *Idaeobatus* | *Idaeobatus* | Asia | 4 | Eurard 11660, MO |
| *R. pentagonus* Wall.<br>Ex Focke | 4x | *Idaeobatus* | *idaeobatus* | Asia | 5 | Alice, Sutherland and Dorji from Bhutan in Alice, Sutherland and Dorji (2008) from Rubus-Ribes 2005. Vouchered WKU 04-06 |
| *R. thomsonii* Focke | 4x | *Idaeobatus* | *idaeobatus* | Asia | 5 | Alice, Sutherland and Dorji from Bhutan in Alice, Sutherland and Dorji (2008) from Rubus-Ribes 2005. Vouchered WKU 04-31 |
| *R. alexeterius* Focke | 2x | *Idaeobatus* | *Idaeobatus* | Asia | 7 | Alice, Sutherland and Dorji from Bhutan in Alice, Sutherland and Dorji (2008) from Rubus-Ribes 2005. Vouchered WKU 04-23 |
| *R. coreanus* Miq. | 2x | *Idaeobatus* | *Idaeobatus* | Asia | 7 | PI 618447/CRUB 1438.001 PL |
| *R. idaeus* L. | 2x | *Idaeobatus* | *Idaeobatus* | Europe/Asia | 7 | T. Eriksson 735, S |
| *R. innominatus* S. Moore | 2x | *Idaeobatus* | *Idaeobatus* | Asia | 7 | PI 553646/CRUB 1039.001 PL |
| *R. lasiostylus* Focke | 2x | *Idaeobatus* | *Idaeobatus* | Asia | 7 | PI 553668/CRUB 425.001 PL |
| *R. leucodermis*<br>Douglas ex Torr. & A. Gray | 2x | *Idaeobatus* | *Idaeobatus* | North America | 7 | PI 553673/CRUB 14.001 PL |
| *R. niveus* Thunb. | 2x | *Idaeobatus* | *Idaeobatus* | Asia | 7 | PI 553723/CRUB<br>269.001 PL |
| *R. occidentalis* L. | 2x | *Idaeobatus* | *Idaeobatus* | North<br>America | 7 | AliceR16,MAINE |
| *R. phoenicolasius*<br>Maxim. | 2x | *Idaeobatus* | *Idaeobatus* | Asia | 7 | Alice96-2,MAINE |
| *R. pungens* Cambess. | 2x | *Idaeobatus* | *Idaeobatus* | Asia | 7 | PI 553849/CRUB<br>46.002 PL |
| *R. sachalinensis* H.<br>Lév. | 4x | *Idaeobatus* | *Idaeobatus* | Asia | 7 | PI 553866/CRUB<br>626.001 PL |
| *R. strigosus* Michx. | 2x | *Idaeobatus* | *Idaeobatus* | North<br>America | 7 | Maine Alice R8 |
| *R. macraei* A. Gray | 6x | *Idaeobatus* | *n/a* | North<br>America<br>(Hawaii) | 6 | Gardners. n., HPDL207 |
| Logan | 6x | *Idaeorubus* | *n/a* | Cultivar | 7 | PI 553258/CRUB<br>81.001 PL |
| Boysen | 7x | *Idaeorubus* | *n/a* | Cultivar | 8 | PI 553341/CRUB<br>1108.001 |
| Marion | 6x | *Idaeorubus* | *n/a* | Cultivar | 8 | PI 553254/CRUB<br>385.001 PL |
| *R. assamensis* Focke | 4x | *Malachobatus* | *Malachobatus* | Asia | 5 | PI 618433/CRUB<br>1701.001 PL |
| *R. ichangensis* Hemsl.<br>& Kuntze | 4x | *Malachobatus* | *Malachobatus* | Asia | 5 | PI 618453/CRUB<br>1606.001 PL |
| *R. irenaeus* Focke | 6x | *Malachobatus* | *Malachobatus* | Asia | 5 | PI 618550/CRUB<br>1607.001 PL |
| *R. lambertianus* Ser. | 4x | *Malachobatus* | *Malachobatus* | Asia | 5 | Boufford &<br>Bartholomew 23955,<br>MO |
| *R. lineatus* Reinw. | 4x | *Malachobatus* | *Malachobatus* | Asia | 5 | Grierson & Long 1950, GH |
| *R. clinocephalus* Focke | 4x | *Malachobatus* | *Malachobatus* | Asia | 5 | PI 606459/CRUB 1642.001 PL |
| *R. tephrodes* Hance | 4x | *Malachobatus* | *Malachobatus* | Asia | 5 | Yao 9231, MO |
| *R. australis* G. Forst. | 4x | *Micranthobatus* | *Lampobatus* | New Zealand | 8 | Gardner 1539, MO |
| *R. parvus* Buchanan | 4x | *Micranthobatus* | *Lampobatus* | New Zealand | 8 | Alice 97-3, MAINE |
| *R. moorei* F. Muell. | 4x | *Micranthobatus* * | *Lampobatus* | Australia | 8 | Streimann 8207, GH |
| *R. calophyllus* | 4x | *n/a* | *Malachobatus* | Asia | 5 | Alice, Sutherland and Dorji from Bhutan in Alice, Sutherland and Dorji (2008) from Rubus-Ribes 2005. Vouchered WKU 04-24 |
| *R. repens* (L.) Kuntze | 2x | *n/a* | *Dalibarda* | North America | 1 | Alice 97-4, MAINE |
| *R. ursinus* × *R. armeniacus*(1) | 8x | *n/a* | *n/a* | North America | 8 | Alice personal collection |
| *R. ursinus* × *R. armeniacus*(6) | 8x | *n/a* | *n/a* | North America | 8 | Alice personal collection |
| *R. acanthophyllus* Focke | 6x | *n/a* | *Orobatus* | South America | 8 | Alice and Cantrell are collectors in Ecuador WKU 07-11 |
| *W. fragarioides* (Michx.) Tratt. | 2x | n/a | n/a | North America | Outgroup | Hill & Soblo 21384, GH |
| *R. glabratus* Kunth | 6x | *Orobatus* | *Orobatus* | South America | 8 | PI 548901/CRUB 1251.004 PL |
| *R. loxensis* Benth. | 6x | *Orobatus* | *Orobatus* | South America | 8 | Alice and Cantrell are collectors in Ecuador WKU 07-17 |
| *R. roseus* Poir. | 6x | *Orobatus* | *Orobatus* | South America | 8 | Luteyn & Quezada 14402, MO |
| *R. laegaardii* Romol. | 6x | *Orobatus* * | n/a | South America | 8 | Alice and Cantrell are collectors in Ecuador Voucher WKU 07-15 |
| *R. hispidus* L. * | 2x | *Rubus* | *Eubatus* | North America | n/a | Alice R9, MAINE |
| *R. caesius* L. | 4x | *Rubus* | *Eubatus* | Europe/Asia | 6 | Karlen 243, S |
| *R. ursinus* Cham. Et. Schltdl. (2) | 8x | *Rubus* | *Eubatus* | North America | 6 | PI 604641 CRUB 1857.001 PL |
| *R. ursinus*(3) | 12x | *Rubus* | *Eubatus* | North America | 6 | PI 554067/CRUB 197.001 PL |
| *R. ursinus*(4) | 13x | *Rubus* | *Eubatus* | North America | 6 | USDA Accession no longer exists |
| *R. ursinus*(5) | 6x | *Rubus* | *Eubatus* | North America | 6 | PI 604641/CRUB 1857.001 PL |
| *R. allegheniensis* Porter | 2x | *Rubus* | *Eubatus* | North America | 8 | Alice R1, MAINE |
| *R. argutus* Link | 2x | *Rubus* | *Eubatus* | North America | 8 | Alice & Judd 15, MAINE |
| *R. armeniacus* Focke | 4x | *Rubus* | *Eubatus* | Europe/Asia | 8 | PI 618579/CRUB 45.001 PL |
| *R. bifrons* Vest | 4x | *Rubus* | *Eubatus* | Europe/Asia | 8 | Alice 98-9, MAINE |
| *R. canadensis* L. | 2x | *Rubus* | *Eubatus* | North America | 8 | Alice & Campbell 98-10, MAINE |
| *R. caucasicus* Focke | 4x | *Rubus* | *Eubatus* | Europe/Asia | 8 | PI 553143/CRUB 54.001 PL |
| *R. coriifolius* Liebm. | 2x | *Rubus* | *Eubatus* | Americas | 8 | Alice and Dodson from Mexico. Vouchered WKU 06-05 |
| *R. cuneifolius* Pursh | 2x | *Rubus* | *Eubatus* | North America | 8 | Alice 5, MAINE |
| *R. flagellaris* Willd. | 4-9x | *Rubus* | *Eubatus* | North America | 8 | PI 553787/CRUB 61.001 PL |
| *R. laciniatus* Willd. | 4x | *Rubus* | *Eubatus* | Europe/Asia | 8 | PI 618548/CRUB 1596.001 PL |
| *R. robustus* C. Presl | 2x | *Rubus* | *Eubatus* | Americas | 8 | Steinbach 247, GH |
| *R. setosus* Bigelow | 2x | *Rubus* | *Eubatus* | North America | 8 | Alice 113, MAINE |
| *R. trivialis* Michx. | 2x | *Rubus* | *Eubatus* | North America | 8 | Alice 33, MAINE |
| *R. ulmifolius* Schott | 2x | *Rubus* | *Eubatus* | Europe/Asia | 8 | 190-84, MOR |
| *R. urticifolius* Poir. | 2x | *Rubus* | *Eubatus* | Americas | 8 | PI 548929/CRUB 1288.001 PL |
| *R. glaucus* Benth. | 4x | *Rubus* | *Idaeobatus* | South America | 6 | PI 548906/CRUB 1293.001 PL |
| *R. eriocarpus* Liebm. | 2x | *Rubus* | *Idaeobatus* | South America | 7 | Alice and Dodson from Mexico. Vouchered WKU 06-12 |
| *R. pensilvanicus* Poir. | 4x | *Rubus* | n/a | North America | 8 | Alice R5, MAINE |

Subg. *Rubus* occurs in the Americas and Europe while *Idaeobatus* is distributed in North America, Europe and Asia; *Malachobatus* is Asian (1, 9–11). *Micranthobatus* and *Lampobatus* were sect. in Focke for species from Australia, Tasmania, and New Zealand (13, 14). Some subg. *Dalibarda* species were moved to subg. *Cylactis* (15). The Flora of China (16), which did not consider global taxa, regrouped species into eight sections corresponding to Focke’s subgenera of similar names. China is a center of species diversity with 139 endemics (12).

Alice and Campbell (17) published a molecular phylogenetic study that sampled the 12 classic subgenera and found that *Orobatus* and *Rubus* were the only monophyletic subgenera. Three major clades were strongly supported. That study underscored the need for additional molecular data to better resolve species level relationships, particularly for polyploids. Asian *Rubus* species were examined using limited nuclear and chloroplast loci by Wang et al. (18). Species from *Dalibardastrum* and *Idaeobatus* were nested within the paraphyletic *Malachobatus*. These authors hypothesized that the allopolyploid species in *Malachobatus* may be derived from crosses between *Idaeobatus* and *Cylactis* species (8, 18). Similarly, *Dalibardastrum* species may be derived from *Malachobatus* ancestors. *Idaeobatus* was polyphyletic with members in four clades. Current phylogenies consistently indicate that subgeneric labels rarely represent monophyletic groups (17, 18).

Hybridization and polyploidization are major evolutionary forces in *Rubus.* Intraspecific morphological and ploidial variability and the capability of many species to hybridize widely across the genus complicate traditional taxonomic classification (8, 19–21). Past phylogenetic analyses of the genus were based on nuclear ribosomal DNA internal transcribed spacer (ITS) sequence data and a few other nuclear and chloroplast loci, including *GBSSI-2, PEPC, trnL/F, rbcL, rpl20-rps*12, and *trnG-trnS* (17, 18, 22). Relying on a limited number of loci to determine relationships in this genus with prevalent hybridization and polyploidy has resulted in low phylogenetic resolution. Additionally, single gene trees may not represent species trees due to hybridization, incomplete lineage sorting (ILS), and gene duplication (23).

Two contrasting views of *Rubus* evolution exist. One view uses a nuclear ribosomal ITS-based genus-wide phylogeny (17) to suggest that the ancestral area for the genus was North America, Eastern Europe (possibly Russia) or Asia (possibly Korea or Japan). In contrast, the treatment of Chinese *Rubus* by Lu (24) hypothesizes that China, where *Rubus* is highly diverse, is the origin of the genus.

In an analysis of Rosaceae using 19 fossils, 148 species and hundreds of low copy nuclear loci, Xiang et al. (25) estimated that this genus originated in the Late Cretaceous approximately 75 Ma. Zhang et al. (26) estimated the age of the root node in a family-wide study of plastid sequences to be 57-66 Ma. *Rubus* fossils exist from the Tertiary period in the Eocene, which began ∼55 Ma, and the more recent Oligocene, Miocene and Pliocene ages, on both sides of the North American land bridge and the Bering land bridge (27).

Certain biogeographical aspects are important to consider for *Rubus* evolution. The North American land bridge connected eastern North America with Europe and Asia before breaking up ∼30 Ma, while the Bering land bridge remained intact until ∼5 Ma (28, 29). Both of these land bridges were important distribution avenues for subtropical (during the warmer Eocene) and temperate species throughout the Tertiary period (28, 30–32). The Panamanian Isthmus connecting Central and South America began closing during the Paleogene approximately 30 Ma. It was crossable for plants and animals at approximately 20 Ma before finally closing 3 Ma (33).

Target capture allows hundreds to thousands of targeted loci to be sequenced for multiple individuals efficiently within a single high-throughput sequencing using Illumina® (San Diego, CA) lane. This technique has resolved phylogenetic questions across a range of plant genera, including *Asclepias* L. (34), *Heuchera* L. (35), and *Lachemilla* L. (36). Although not specifically targeted, chloroplast sequences can be obtained after sequencing target capture libraries, enabling an independent estimate of phylogeny and inference from a predominantly maternally inherited genome (34, 35, 37).

The objectives of this study were to estimate phylogenetic relationships in *Rubus* using a large molecular dataset over a genus-wide species sampling; estimate divergence times between major *Rubus* clades; and examine the biogeography of species diversification.

## Results

### Sequencing target genes and chloroplast genome

The average sequencing depth for all samples over all loci was 66.8x (Additional file 1). The samples of *Rubus hispidus* and *R. pectinellus* had an average sequencing depth across all loci under 1x and contigs for <10 genes and were excluded from phylogenetic analyses. The average percentage of on-target reads was 71.3%. HybPiper produced sequences for an average of 1,113 genes per taxon, and an average of 988 sequences were at least 75% of the target length. An average of 86% of target bases were recovered for genes shared across Rosaceae and 101% of bases for *R. occidentalis* targets (Additional file 2). Alignment lengths for supercontigs, *i*.*e*., exons + noncoding sequences, were 10.1 Mbp for diploid species only (average ungapped length 3.8 Mbp) and 10.1 Mbp for polyploid and diploid taxa (average ungapped length 2.7 Mbp (Additional file 2). The concatenated alignment length of exon sequences for each gene was 2.5 Mbp for diploid species only (average ungapped length 1.6 Mbp) and 2.5 Mbp for all analyzed taxa (average ungapped length 1.7 Mbp). The supercontig sequence alignments of diploids and all species, had 17% and 23% variable sites and 7% and 11% phylogenetically informative sites, respectively. Exon alignments were 20% variable (9% phylogenetically informative) for diploids and 29% variable (15% informative) for all species analyzed.

After automated trimming and manual evaluation of alignment quality, 941 gene targets remained for exon alignments and 905 to 910 for supercontigs from all taxa, and from diploids only, respectively (38). After removal of rogue taxa (those with ambiguous phylogenetic placement), exon alignments of all taxa and alignments of only diploid taxa, contained an average of 52 (55% of total sample set) and 30 taxa (70% of total sample set), respectively. Supercontig alignments including all taxa contained an average of 39 taxa (41% of total sample set), while alignments of only diploid taxa contained 27 individuals on average, or 63% of total sample set (38).

The chloroplast alignment of sequences from 89 taxa was 125,795 bp. RogueNaRok identified *R. caucasicus, R. lambertianus* and *R. robustus* as rogue taxa and they were removed from the chloroplast analysis. Average coverage of the 127,679 bp *R. occidentalis* reference genome was 24x, ranging from 1.3x-99.6x (Additional file 3).

### Phylogenetic analyses

Differences between the ASTRAL-II and SVDQuartets analyses for all taxa and diploid-only taxa datasets were more evident in the topology of internal nodes delineating the relationships between groups (Fig. 1). These nodes represent relationships between groups that may commonly hybridize or where ancestors of extant taxa may have been progenitors of multiple clades. Deep evolutionary signal for these events may have been obscured by more recent polyploidization and hybridization events, leading to topological conflict between analyses. The quartet concordance (QC) values for two nodes describing relationships between major groups in the SVDQuartets phylogenies indicate counter support for the topology. The alternate topologies seen in the ASTRAL-II trees have weak support and skewed distributions for discordant topology frequencies at some internal nodes (38). The SVDQuartets trees are likely exhibiting these discordant topologies that are supported by a significant minority of loci. In a previous report, ASTRAL-II phylogenies were shown to be more accurate than SVDQuartets trees in the presence of high ILS (39).

**Figure 1.**
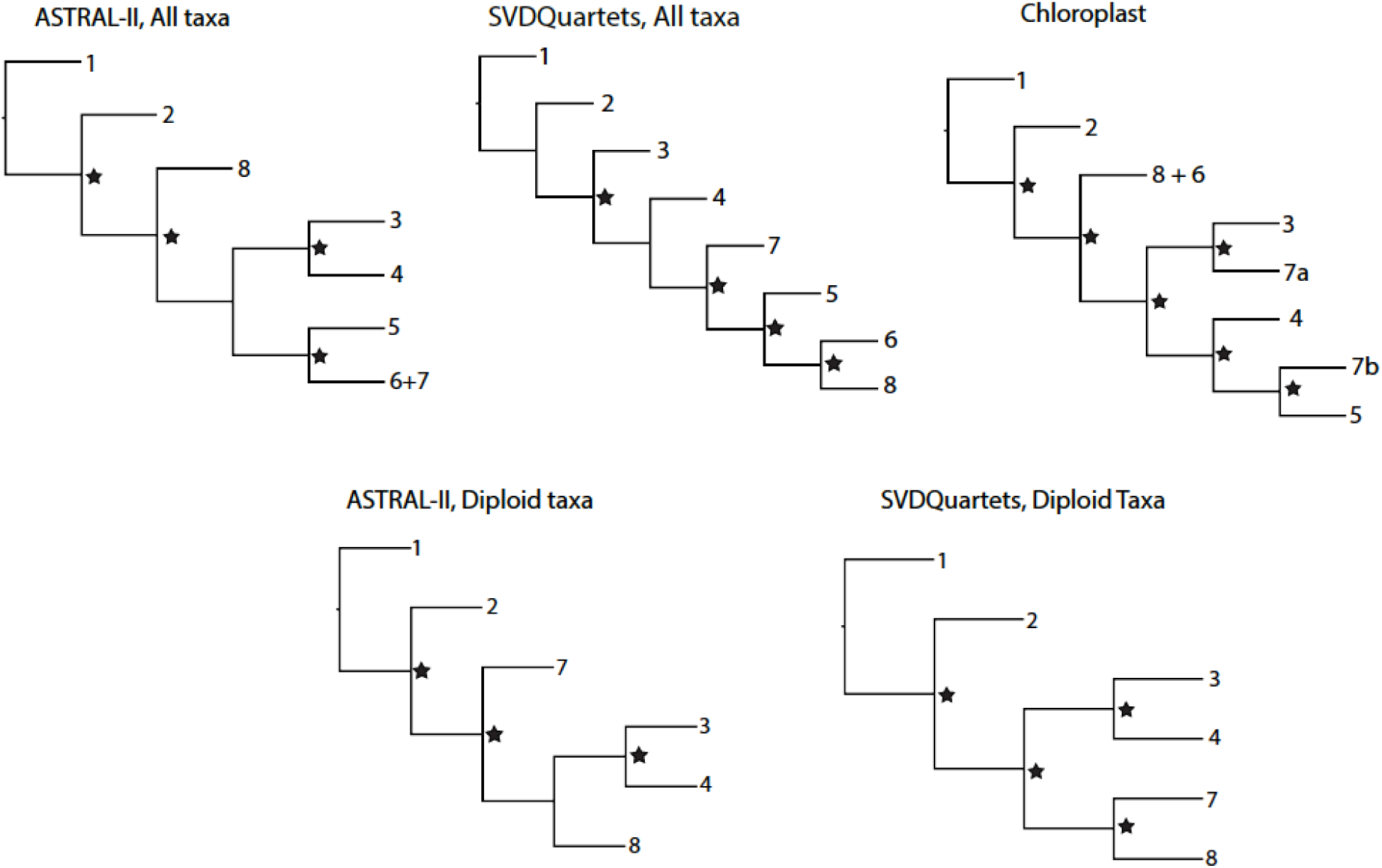
Topological relationships between genus *Rubus* Groups 1-8 in phylogenetic analyses of exon or chloroplast sequences. Nodes with strong support (Bootstrap > 75 for SVDQuartets hylogenies; Posterior Probability > 0.95 for ASTRAL-II phylogenies) are marked with a star.

The supercontig sequence alignments contained a high proportion of missing data. On average, 73% of the data was missing from the supercontig sequence alignments for all taxa, compared to 42% of missing data for the exon alignment for all taxa (38). Similarly, diploid alignments had an average of 64% missing data for supercontig sequence data and 39% for exon sequences. When compared, the exon-only phylogenies and the supercontig sequence trees show the same major groups of taxa and similar variations in backbone topologies between analyses (Fig. 1; exon-only phylogenies). Because the supercontig sequence dataset did not provide additional phylogenetic resolution and contained less complete alignments, we focused on analyses of the exon sequences.

Eight consistent groups of taxa corresponding roughly to eight clades were seen in the SVDQuartets and ASTRAL-II generated phylogenies from all datasets: diploid exons, diploid supercontig sequences, polyploid and diploid exons, and polyploid and diploid supercontig sequences (Fig. 2, 3,). Most relationships in the analyses were well-supported (bootstrap values > 75; posterior probabilities > 0.95). In addition, group 8 was divided into 8a representing the majority of this clade and subg. *Rubus*; group 8b, including subg. *Orobatus* species; and group 8c, including subg. *Comaropsis, Diemenicus,* and *Micranthobatus*. Most relationships in the analyses were well-supported (bootstrap values > 75; posterior probabilities > 0.95).

**Figure 2.**
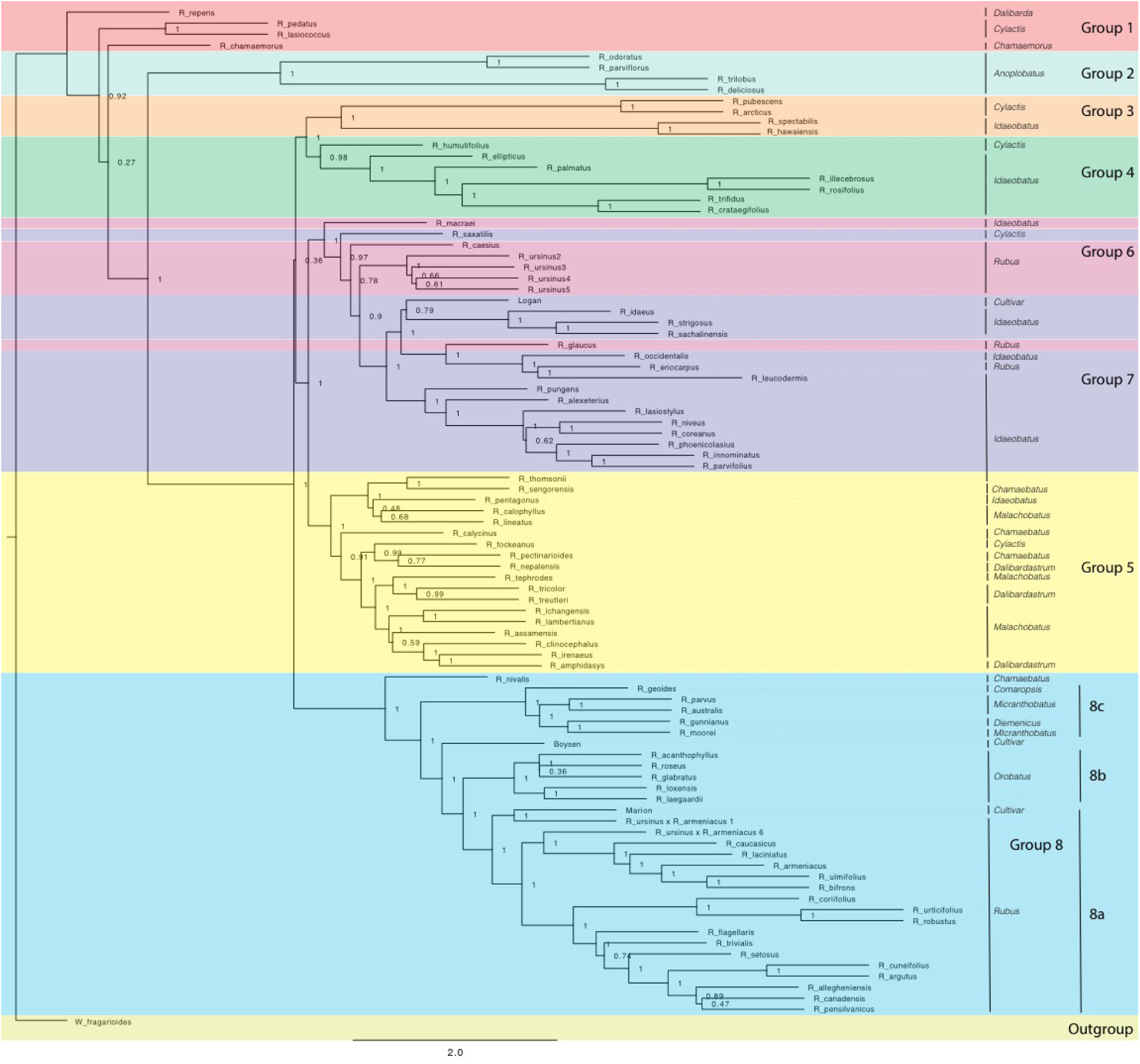
ASTRAL-II phylogeny estimated from exon sequence gene trees from all *Rubus* taxa. Posterior probability values (0–1) are shown to the right of each node. Branch lengths are in coalescent units and measure discordance in the underlying gene trees. Groups are labelled with colored bands. Taxa are labelled with their subgeneric classification.

Groups 1 and 2 include eight species from subg. *Chamaemorus*, *Dalibarda*, *Cylactis*, and *Anoplobatus* and are sister to the remainder of genus *Rubus* (Fig. 2, Table 1). Group 3 includes *R. hawaiensis* (*Idaeobatus*), *R. spectabilis* (*Idaeobatus*), *R. pubescens* (*Cylactis*), and *R. arcticus* (*Cylactis*), and is monophyletic. Group 4 is sister to Group 3 and contains seven taxa; six are classified in *Idaeobatus* and one in *Cylactis* (*R. humulifolius*). Group 5 consists of accessions of Asian origins from *Malachobatus*, *Daliardastrum*, *Cylactis*, *Idaeobatus*, and *Chamaebatus*. It is often sister to Group 7, which contains primarily *Idaeobatus* species with one *Cylactis* accession (*R. saxatilis*) and ‘Logan’, a hybrid cultivar. Group 6, contains four of six *R. ursinus* accessions, *R. caesius*, and *R. glaucus* from subg. *Rubus* and *R. macraei* from *Idaeobatus,* and shifts positions between analyses but groups with either Group 7 or 8. Group 8 contains the most species and consists of accessions from subg. *Rubus* (8a), *Orobatus* (Group 8b), *Comaropsis*, *Micranthobatus*, and *Diemenicus* (8c), and the predominantly blackberry hybrid cultivars ‘Boysen’ (75% blackberry/25% raspberry) and ‘Marion’ (69% blackberry and 31% raspberry). *Anoplobatus* and *Orobatus* are monophyletic (Fig. 2). All other subgenera, except monotypic *Chamaemorus, Comaropsis* and *Diemenicus*, are para- or polyphyletic. *Anoplobatus* species comprise Group 2 and are sister to the majority of genus *Rubus. Orobatus* species form a subclade in Group 8 and are sister to the major subg. *Rubus* clade. Species from *Comaropsis, Micranthobatus, and Diemenicus* also form a subclade in Group 8. Subg. *Rubus* would be monophyletic in Group 8 if not for *R. ursinus, R. glaucus,* and *R. caesius* in the variable Group 6, and *R. eriocarpus* in Group 7. These species are putative allopolyploids and are discussed below. Species from *Comaropsis, Micranthobatus, and Diemenicus* form a subclade in Group 8. In the chloroplast phylogeny, Group 7 divides into two monophyletic clades. One is sister to Group 3 and the other to Group 5. The eight major groups also appear in phylogenetic network analyses (Fig. 4, 5).

**Figure 3.**
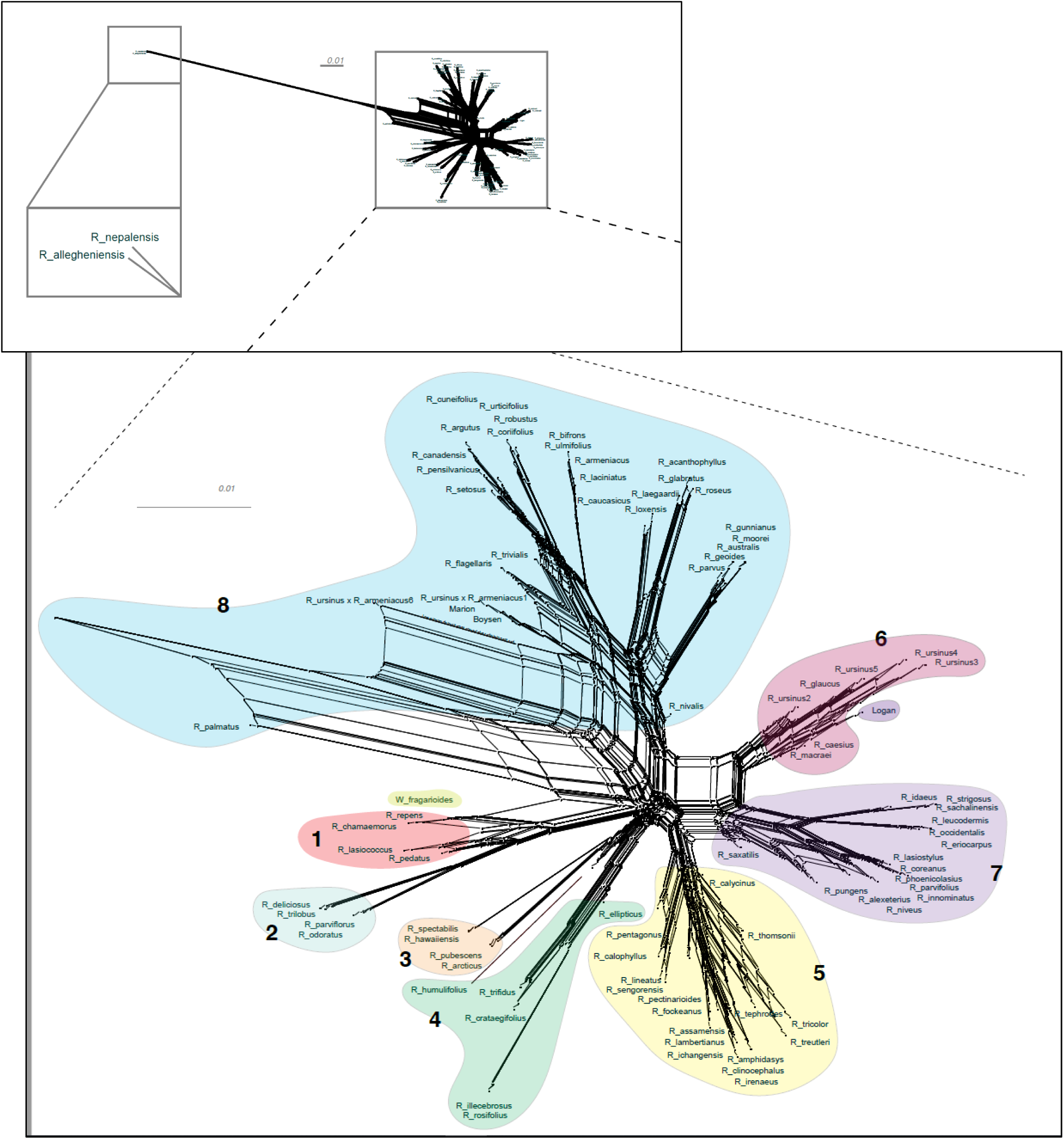
Super network for all *Rubus* taxa estimated with SuperQ from exon gene trees estimated with RAxML. Colored shapes correspond to Groups 1-8. Top inset placement of R. allegheniensis and R. nepalensis due to limited sequence data for these samples (38).

**Figure 4.**
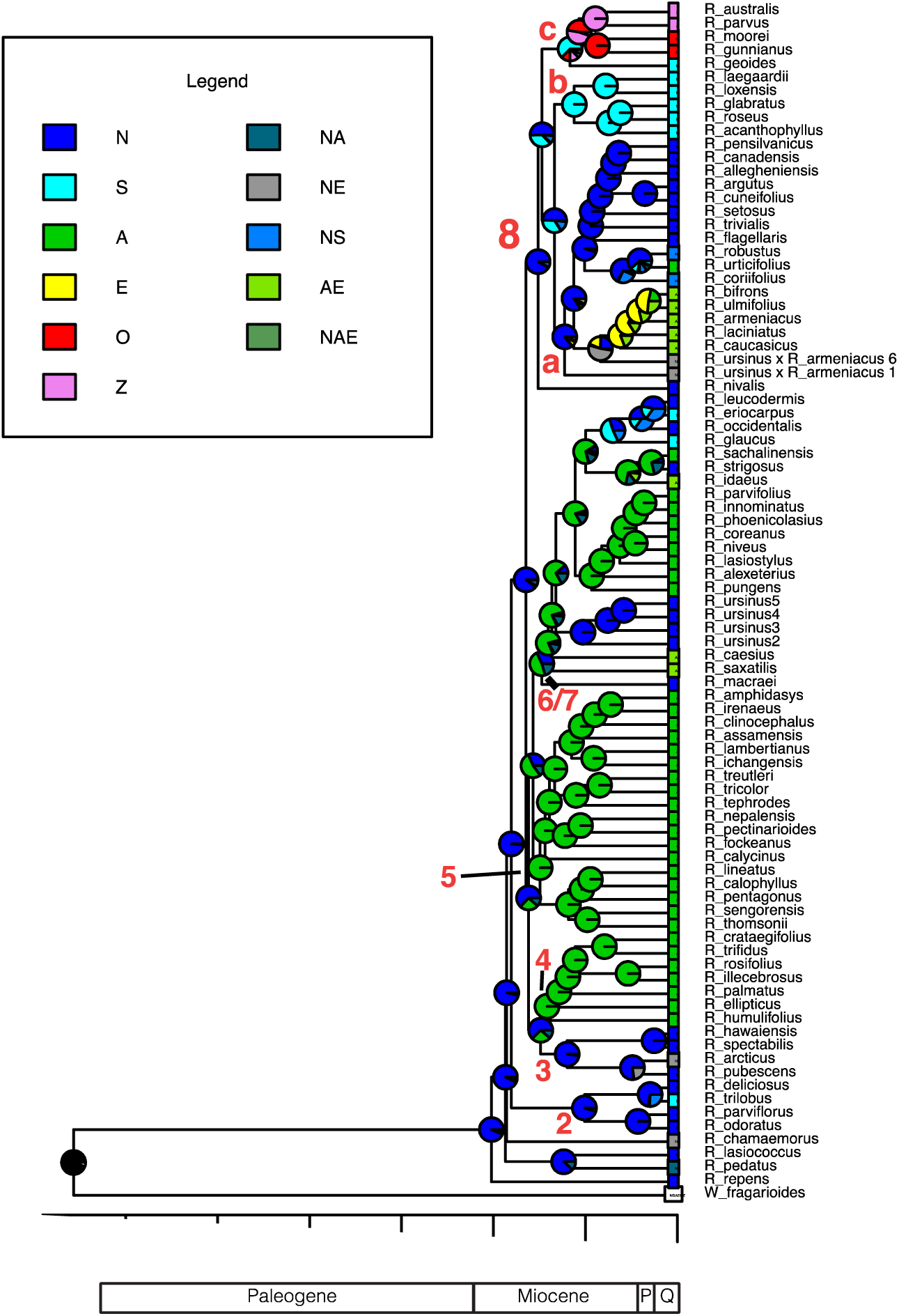
*Rubus* ancestral range estimation using the DEC+*j* model for all taxa. Time scale is in millions of years. Pie charts represent relative probability of each area being the ancestral range. P=Pliocene, Q=Quaternary, N=North America (including Mexico and Guatemala), S=South America, A=Asia, E=Europe, O=Australia, Z=New Zealand. Combinations of letters indicate presence across multiple areas. Ancestral nodes for major groups are labelled numerically.

Network analyses allowed a more thorough visualization of conflict within our data, particularly caused by hybridization, as discussed below, which cannot be captured in a dichotomously branching tree (Fig. 3).

Maternal and paternal progenitors of putative hybrid groups or species were assessed by comparing nuclear and chloroplast phylogenies (Fig. 2, 3). *R. nepalensis* and *R. allegheniensis* had long branch lengths compared to other taxa (Fig. 3), likely due to limited sequence data for these samples (38). These species have sequences over 75% of the target length for only 89 and 66 targets, respectively. ASTRAL-II and SVDQuartets are robust to this level of missing data and place these species with high support in most species trees (Fig. 2).

### Phylogenetic dating and ancestral range estimation

Ultrametric trees of all taxa estimated from exon sequences and dated using r8s are shown (Fig. 4). *Rubus* radiated throughout the Miocene with the eight major groups arising approximately 10-20 Ma. The DEC model for ancestral range estimation was rejected based on a likelihood ratio test (*p*<0.05) and AIC values (38). Under the DEC+*j* model, the most likely ancestral range for *Rubus* for all taxa phylogenies was North America (Fig. 4). Most recent common ancestors (MRCA) of Groups 1, 2, 3, and 8 were also most likely distributed in North America. Ancestral ranges in North America and Asia were similarly likely for Group 6+7 (Fig. 4). Ancestors of Groups 4 and 5 were most likely distributed in Asia.

## Discussion

### Phylogenetic analyses and taxonomic implications

Our target capture sequencing approach enabled resolution of relationships between major groups, confirming or extending from previous studies (17, 18, 22, 40). Subg. *Idaeobatus* is polyphyletic, as seen in studies of Asian and worldwide *Rubus* species (17, 18, 22, 40). *Rubus macraei* and *R. hawaiensis* have distinct evolutionary histories and likely resulted from separate colonization events of the Hawaiian Islands (40). *Rubus ursinus* is not closely related to *R. macraei* (Fig. 3).

*Rubus repens* is placed in genus *Rubus* as *R. dalibarda* (9–11), but classified by other botanists in the monotypic genus *Dalibarda* due to unique morphological features rarely or not otherwise seen in *Rubus,* including dry fruits, reduced carpel number, and apetalous, dioecious flowers (15, 17, 41). Alice and Campbell (17) showed this species nesting within subg. *Rubus* using ITS data, spurring its reclassification into genus *Rubus* (21). In our study, *R. repens* nests either within Group 1 or is sister to other *Rubus* species studied, supporting its classification in the genus *Rubus*.

The six subg. *Cylactis* species are distributed in Groups 1, 3, 4, 5, and 7 and often closely related to species in subg. *Chamaebatus* or *Idaeobatus* (Fig. 2). Morphological differences used for current taxonomic classifications in Group 5 do not reflect phylogenetic relationships. Since many of these species are likely allopolyploids with similar progenitor species, taxonomy based on morphology may be unreliable for this group.

Subgenus *Micranthobatus* is closely related to the monotypic subg. *Diemenicus* and *Comaropsis.* Several species in these subgenera are tetraploid with small genomes (42). Our results support the hypothesis of Hummer and Alice (42) that these species may have descended from one allopolyploid ancestor, possibly a hybrid between diploids with small genomes. *Rubus nivalis,* a closely related diploid species, may have been a progenitor of this group. The common ancestor of these five species may have migrated from South America to the South Pacific through long distance dispersal by birds. Geographic isolation, potentially between populations of the common ancestor of *R. moorei* and *R. gunnianus*, may have led to strong morphological divergence. *Rubus gunnianus* of the monotypic subg. *Diemenicus* has unique morphological features, including leaves arising in rosettes directly from the rhizome, a lack of stipules, broad petioles, prominent carpel glands, and unisexual flowers (14).

Subg. *Rubus* species are primarily in Groups 6 and 8, with *R. eriocarpus* in Group 7. *Rubus eriocarpus* is morphologically similar to *R. glaucus* (43). Both share stem and leaf characteristics with black raspberries but have fruit that retains the torus like a blackberry when picked (3, 44). *Rubus eriocarpus* is closely related to North American black raspberries *R. occidentalis* and *R. leucodermis* in nuclear and chloroplast phylogenies (Fig. 2, 5) while *R. glaucus* aligns with other putative blackberry × raspberry hybrids in Group 6. Focke (9–11) originally classified *R. eriocarpus* in *Idaeobatus*; our results support Focke’s treatment of *R. eriocarpus* within subg. *Idaeobatus*. Similarities between *R. glaucus* and *R. eriocarpus* could be due to convergent evolution, or *R. eriocarpus* could be a parent of *R. glaucus*.

Subgenus *Idaeobatus* is polyphyletic with representatives in Groups 3, 4, 5, and 7. Groups 7 and 4 contain primarily *Idaeobatus* species, but they are not closely related. Group 4 is highly supported as sister to Group 3 in analyses of exon sequences for all taxa (Fig. 2) as well as for diploid taxa only (38). Group 7 further splits into two separate groups in the chloroplast analysis (Fig. 5). One branch is sister to Group 3 while the other is sister to Group 5, indicating strong maternal genetic differences between these two *Idaeobatus* groups. Multiple studies have recognized polyphyly in *Idaeobatus* (17, 18, 22, 40). High support for divisions between *Idaeobatus* species in this and other studies indicate that this subgenus would benefit from further phylogenetic study and taxonomic reclassification.

**Figure 5.**
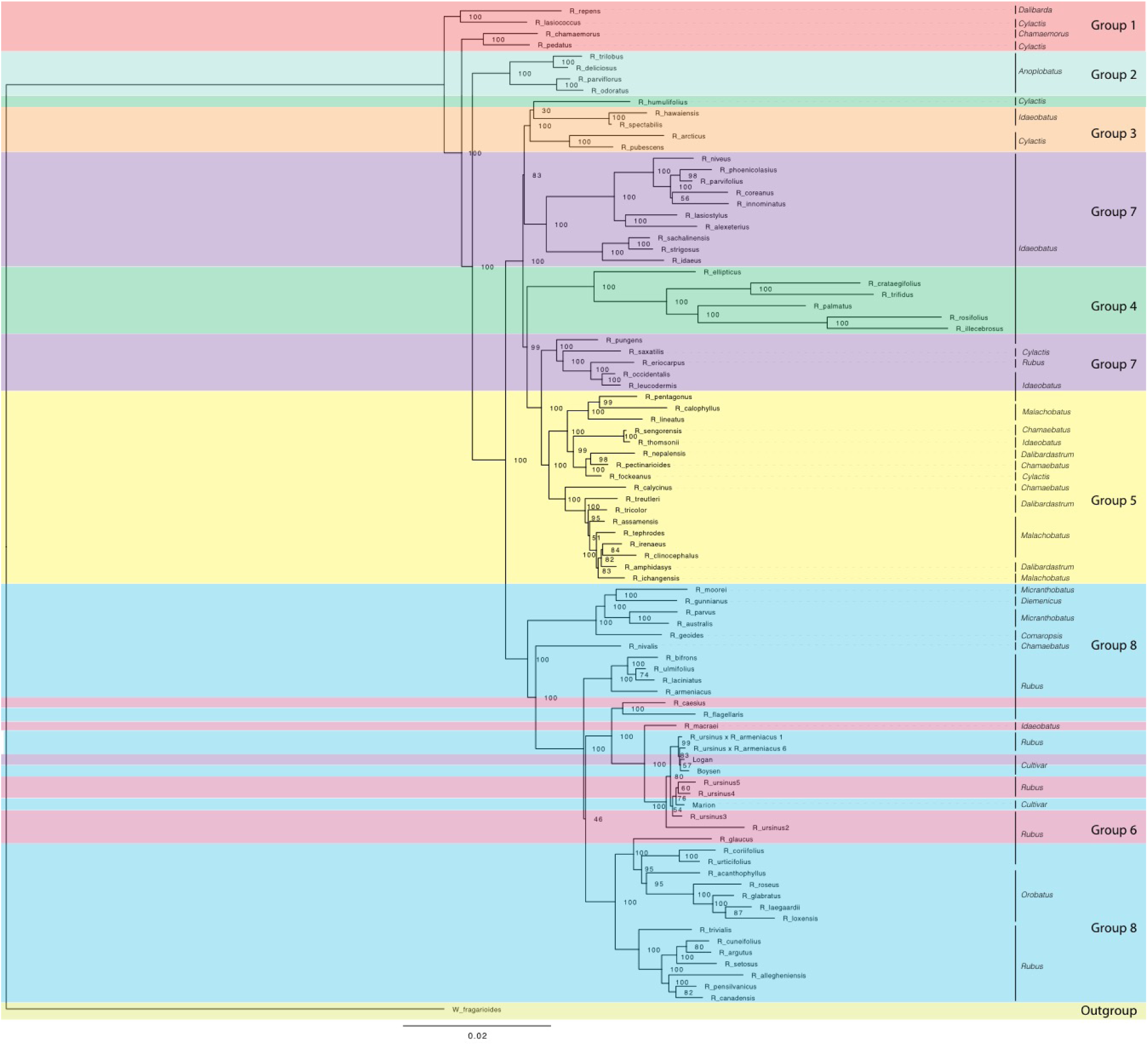
Maximum likelihood phylogeny estimated with RAxML for chloroplast sequences from all *Rubus* taxa. Bootstrap values (0–100) are shown to the right of each node. Branch lengths represent relative evolutionary change. Groups are labelled with colored bands. Taxa are labelled with their subgeneric classification sensu GRIN (2019).

### Hybrids

The HybPiper assembly pipeline reduced the complexity of polyploid species by choosing the longest sequence per target locus (45). Because there are hundreds of targets, the evolutionary history of each subgenome in a polyploid was represented by a proportion of the loci, thus, the species trees give a broad overview of that mixed signal. Dichotomous trees place hybrid taxa intermediately between progenitors because their genomes have conflicting phylogenetic signal (46). However, if parents are distantly related, the hybrid taxon may not appear phylogenetically close. Without the constraint of dichotomous branching, network analyses allowed a more thorough visualization of such conflict within our data and possible hybrids. *Rubus* hybrids ‘Logan’, ‘Boysen’, and ‘Marion’ are horticulturally and economically important cultivars in major berry production regions in the Pacific Northwest and around the world (3, 4, 47). All three are known blackberry × raspberry hybrids. ‘Logan’ has the closest raspberry relative with ‘Red Antwerp’ as the documented pollen parent (3). ‘Boysen’ is an offspring of ‘Logan’ and thus has a raspberry grandparent. ‘Logan’ and ‘Boysen’ are both derived from ‘Aughinbaugh’, a domesticated western North American *R. ursinus* selection (3). ‘Marion’ has a raspberry for a great-great-grandparent and is also related to *R. ursinus* (Thompson 1995, Jennings 1988). All three cultivars cluster with the *R. ursinus* selections in the chloroplast phylogeny, confirming the documented relationships with this species (Fig. 5). In nuclear analyses, ‘Logan’ groups with other raspberries in Group 7 while ‘Boysen’ and ‘Marion’ are positioned in Group 8 with the blackberries (Fig. 2). The position of ‘Logan’ with the raspberries is as expected given its paternal red raspberry parent and the possibility that *R. ursinus* may also be a hybrid berry (17, 40). QC values are low or negative for ‘Boysen’ and ‘Marion’ related nodes, indicating that a weak majority or minority of quartets support the position of these species (38). The raspberry germplasm in their recent heritage creates conflict in the phylogenetic signal for these taxa. In network analyses, ‘Marion’ and ‘Boysen’ group with other blackberries in Group 8 while ‘Logan’ is placed within Group 6, between Groups 7 and 8 (Fig. 4). The placement of ‘Logan’ between Groups 7 (raspberries) and 8 (blackberries) reflects its hybrid heritage.

Evidence of hybridization exists across the *Rubus* phylogeny (Fig. 4, Additional file 4). The position of *R. chamaemorus* (2*n* = 8*x* = 56) (4) in Group 1 has low support in the exon-based ASTRAL-II phylogeny (Fig. 2). In a previous study, two *R. chamaemorus* alleles from GBSSI-1γ appeared either outside of the major *Rubus* clade as sister to *R. lasiococcus* or inside as sister to *R. arcticus* (48). *Rubus chamaemorus* may have progenitors outside of and within the major *Rubus* clade, leading to its variable position. The maternal progenitor for *R. chamaemorus* is likely a lineage outside of the major *Rubus* clade since this species is sister to *R. pedatus* in Group 1 in the chloroplast phylogeny (Fig. 5). This finding supports that *R. chamaemorus* may have an autopolyploid origin (49).

*Rubus humulifolius* is strongly associated with Group 4 in the exon ASTRAL-II phylogeny, but groups (with low support) in Group 3 in the chloroplast tree (Fig. 2, 5). In the exon split network, *R. humulifolius* occupies a short node between Groups 3 and 4 (Fig. 3). This indicates that splits in gene trees do not consistently place this species with either Group 3 or Group 4. *Rubus humulifolius* (2*n* = 4*x* = 28) is the only polyploid taxon in either of these two groups, a trait also indicative of hybrid origin. Progenitors are likely from subg. *Idaeobatus* and/or *Cylactis*.

Similar to *R. humulifolius, R. saxatilis* (2*n* = 4*x* = 28) is another polyploid in a primarily diploid clade. *Rubus saxatilis* is closely related to subg. *Idaeobatus* species in Group 7, although it is currently classified in subg. *Cylactis*. In the chloroplast tree, this species is sister to the black raspberries, *R. occidentalis, R. leucodermis, R. eriocarpus,* and *R. pungens* (Fig. 5). Network analyses from exon sequences place *R. saxatilis* between Groups 5 and 7 with a short branch, exhibiting conflict in the placement of this species (Fig. 3). The supercontigs sequence network places *R. saxatilis* unexpectedly near Group 3 along with *R. caesius* (38). The maternal progenitor of this species is likely from subg. *Idaeobatus,* and may be *R. occidentalis*. The paternal parent is unknown and may be a member of Group 3, 5, 6, or 7.

Group 5 members include the Asian polyploids subg. *Malachobatus, Dalibardastrum, Chamaebatus, Cylactis,* and *Idaeobatus*. The diploid exon ASTRAL-II tree shows that Groups 3 and 4 are more closely related to Group 8 than to Group 7 (38). Members of subg. *Idaeobatus*, such as *R. parvifolius* or *R. pentagonus,* and members of subg. *Cylactis*, such as *R. fockeanus,* may have been progenitors of this likely allopolyploid subgenus (18). *Rubus pentagonus* is closely related to subg. *Malachobatus* species in Group 5, along with other subg. *Idaeobatus* taxa, *R. thomsonii* and *R. sengorensis*. The shift in the relationship between Groups 3, 4, 7, and 8 after the addition of putative allopolyploids in Group 5 lends support to the hypothesis that subg. *Malachobatus* is derived from subg. *Idaeobatus* and *Cylactis* species (18). Phylogenetic signal from Group 5 brought the progenitor species and their relatives from Groups 3, 4 and 7 together in the dichotomous phylogeny. *Rubus pentagonus, R. thomsonii,* and *R. sengorensis* may be progenitors of this group or examples of subg. *Idaeobatus* hybrids. In the chloroplast analysis, these three species are embedded within Group 5 with other subg. *Malachobatus* and *Dalibardastrum* species. Sister to Group 5 is another group of subg. *Idaeobatus* species, *R. pungens, R. saxatilis, R. eriocarpus, R. occidentalis*, and *R. leucodermis* that could be possible progenitor species.

Species from subg. *Dalibardastrum,* another polyphyletic subgenus in Group 5, are also putative allopolyploids with progenitor species either from or similar to those for subg. *Malachobatus.* Network analyses distinctly show Group 5 separating from other groups, but the extensive webbing between taxa illustrates conflict in the dataset for these species. This demonstrates the convoluted evolutionary history between these putative allopolyploids. Group 5 is positioned between Groups 7 and Groups 3 and 4, which include the proposed progenitors from subg. *Idaeobatus* and *Cylactis* (Fig. 3).

Blackberry × raspberry hybrids in Group 6 are primarily classified in subg. *Rubus* but are genetically distinct from other blackberries in Group 8 in nuclear analyses. A hybrid subgenus, such as *Idaeorubus* Holub, initially described for cultivars, may be applicable for these taxa. There are two strongly supported subgroups in Group 8. *Subgenus Orobatus* species form one, while Australasian species in subg. *Diemenicus* and *Micranthobatus*, along with southern South American *R. geoides* from subg. *Comaropsis,* form another (Fig. 2). Both subgroups are distinct from but closely related to the major subg. *Rubus* clade. This could be interpreted in two ways. First, populations of the common ancestor of these species may have become reproductively isolated and subsequently evolved into each of these three major groups. It is difficult to reconcile the varying ploidy levels of all species involved with this scenario. Another hypothesis is that both subgroups have one progenitor within or closely related to subg. *Rubus* and another in a different subgenus, such as *Anoplobatus* for *Orobatus* or *Cylactis* for *Comaropsis*/*Diemenicus*/*Micranthobatus* (3, 42). The maternal parent in either of these hypothesized crosses is from subg. *Rubus* because all three are in Group 8 in the chloroplast phylogeny (Fig. 5).

Group 6 contains additional putative hybrids between subg. *Idaeobatus* and subg. *Rubus.* In nuclear phylogenies, this clade shifts positions but is either associated with Group 7 or 8 (Fig. 2). In the chloroplast phylogeny, these species do not form a clade but all group with subg. *Rubus* in Group 8 (Fig. 5). The exon network for all taxa places Group 6 between Groups 7 and 8 (Fig. 3).

*Rubus glaucus* is morphologically similar to black raspberries (Group 7) with semi-erect, glaucus canes and trifoliate leaves, but has fruit that adheres with the torus like a blackberry (3, 9–11, 44), It is closely related to black raspberries *R. eriocarpus, R. occidentalis* and *R. leucodermis* in the exon ASTRAL-II phylogeny of all taxa (Fig 2). In the chloroplast tree, *Rubus glaucus* shifts into Group 8 where it is related to subg. *Rubus* and *Orobatus* taxa (Fig. 4). If it is a cross between a black raspberry and a blackberry, as its morphology suggests and supported by its variable placement with weak support in nuclear phylogenetic analyses, a black raspberry was likely the paternal donor (3, 9–11).

*Rubus caesius* is a tetraploid blackberry that hybridizes readily with other bramble species (3, 7) and has given rise to many new blackberry varieties in Europe (6). The maternal parent for *R. caesius* was likely in subg. *Rubus* given its position in Group 8 in the chloroplast phylogeny (Fig. 4).

*Rubus macraei* and *R. hawaiensis* are both endemic Hawaiian species, but are evolutionarily separate. *Rubus hawaiensis* is in Group 3 and sister to *R. spectabilis* with strong support in all analyses (Fig. 2, 3). *Rubus macraei,* a hexaploid (2*n* = 6*x* = 42) (40), is a member of Group 6 and another putative blackberry × raspberry hybrid. These results support the hypothesis that *R. hawaiensis* and *R. macraei* arose from separate colonization events of the Hawaiian Islands (40, 50).

*Rubus ursinus* is represented by six accessions. Specimens 1 and 6 are putative *R. ursinus* × *armeniacus* hybrids and are in Group 8 in all nuclear analyses (Fig. 2). In the chloroplast phylogeny, they group with the other *R. ursinus* accessions, indicating that *R. armeniacus* was the pollen parent (Fig. 5). Despite varying ploidy levels, *R. ursinus* accessions 2, 3, 4, and 5 in Group 6 form a clade (Fig. 2). Variability in the placement of *R. ursinus* in nuclear phylogenies indicates that the species is a blackberry × raspberry hybrid with the maternal parent in subg. *Rubus* (Fig. 2) (38). This supports the hypothesis in Alice et al. (1999) that *R. ursinus* is a hybrid, however there is no direct evidence that *R. macraei* is a parent of *R. ursinus.* Rather, both of these species are putative blackberry × raspberry hybrids of unknown origin.

### Ancestral ranges and geographic migrations

The *Rubus* MRCA is most likely from North America, supporting the hypothesis presented by Alice and Campbell (17) based on an ITS phylogeny (Fig. 4). This contradicts hypotheses by Lu (24) and Kalkman (51) that *Rubus* originated in southwestern China or Gondwanaland. For *Rubus,* high diversity seen in Asian regions does not correspond with the most likely ancestral range. *Rubus* in Groups 4, 5, 6+7, and 8 colonized Asia at least three times during the Miocene (Fig. 4). Group 5 is likely the result of a hybridization event between progenitors already distributed in Asia since these species are not present in North America. Groups 4, 7 (both primarily subg. *Idaeobatus*) and 8a (primarily subg. *Rubus*), show classic eastern Asian-eastern North American biogeographic disjunction patterns where closely related species are dispersed across both geographic locations (27, 52). During the Miocene, plant dispersal from North America to Asia could have occurred over the Bering or North American Land bridges (28–31). Distributions in Groups 4 and 7 likely occurred over the Bering land bridge because North American species in these groups are presently distributed in western regions. *Rubus sachalinensis*, an Asian red raspberry, is native to Europe and Asia, but clusters with other North American subg. *Idaeobatus* species, including *R. strigosus*, and the European *R. idaeus.* These European species have a unique evolutionary path compared to other Asian subg. *Idaeobatus* taxa and may be another example of an independent *Idaeobatus* migration from North America into Eurasia. This supports results from Wang (53) using *matK* chloroplast sequences to study *Rubus* species used in traditional Chinese medicine where *R. sachalinensis* was sister to *Idaeobatus* accessions from Asia. Morphological stasis may explain why character states do not differentiate these genetically differentiated *Idaeobatus* groups. Stasis occurs when evolutionary constraints and stabilizing selection prevent significant changes in morphological characters between lineages (31, 54). This can occur when disjunct geographic areas have similar habitats, such as in North America and eastern Asia (55).

In Group 8, the Eurasian distribution of many species and the presence of close genetic relatives in eastern North America suggest migration across the North Atlantic land bridge, however this passage closed at the latest 15 Ma (56). North American ancestors of Group 8 taxa may have been widespread across North America in the broadleaved, deciduous, temperate forests characterizing the Miocene (57). These species could have migrated across the Bering land bridge through Asia and into Europe. During the subsequent Pleistocene glaciation events, North American distributions shrank back into the east. After diploid species migrated to Europe through the late Miocene, glacial cycles created conditions beneficial for the success of apomictic polyploids. With populations fragmented among glacial refugia, the ability to reproduce asexually may have been advantageous (6).

Ancestors of species distributed in Mexico, Guatemala, and South America, in Groups 2 and 7 (*R. trilobus, R. glaucus,* and *R. eriocarpus*) may have diversified from their North American relatives. This would have occurred after temperature decreases and the spread of grasslands during the Pliocene created refugia of the widespread broadleaved, deciduous forests of the Miocene in the southeastern US and Mexico (57). In the mid-Miocene, the South American subgenus *Orobatus* diverged from other Group 8 taxa. During the Paleogene, approximately 30 Ma, the Panamanian Isthmus connecting Central and South America began to close. The isthmus was crossable for plants and animals at approximately 20 Ma until 3 Ma (33). *Rubus geoides* in Group 8c also differentiated from North American ancestors during this time frame. Long distance dispersal most clearly explains the disjunction between *R. geoides* in South America and subg. *Micranthobaus/Diemenicus* species in Australia and New Zealand. This vicariance occurs too late (approx. 10 Ma) to have occurred over the land bridge between South America, Antarctica, and Australia, which broke up in the late Eocene approximately 30 Ma, when the continental shelves were no longer exposed (58). A similar dispersal event occurred in Vitaceae and was likely driven by birds (52). Further geographic isolation after dispersal between Tasmania and New Zealand likely led to speciation between *R. parvus* and *R. australis* (New Zealand) and *R. gunnianus* and *R. moorei* (Tasmania) (42).

## Conclusions

*Rubus* phylogenetic estimation has been hindered by a lack of single copy genes because of whole genome duplication and hybridization, resulting in polyploidy. This target capture dataset of approximately 1,000 single copy loci provided high resolution between species for many clades but also evidence of gene tree/species tree and cytonuclear discordance. In most cases, discordance is due to biological processes such as hybridization and incomplete lineage sorting as opposed to a lack of phylogenetic signal (38). This study illustrates the importance of using multiple phylogenetic methods when examining complex groups and the utility of software programs that estimate signal conflict within datasets. Within each clade, taxon composition and relationships were highly consistent. Differences between datasets and analyses were more evident in the topology of internal nodes delineating the relationships between groups where phylogenetic signal may be obscured by recent polyploidization and hybridization events.

*Anoplobatus* and *Orobatus* are the only strictly monophyletic subgenera. Putative allopolyploid subgenera *Dalibardastrum* and *Malachobatus* are closely related and may have progenitors in subg. *Idaeobatus* or *Cylactis*. Subgenus *Idaeobatus* is strongly polyphyletic in nuclear and chloroplast analyses. Subgenus *Rubus* is monophyletic with the exception of putative allopolyploids *R. glaucus, R. caesius, and R. ursinus*.

The analysis of cultivated blackberry × raspberry hybrids with known pedigrees confirms the effectiveness of target capture sequencing for phylogenetic analysis. This approach successfully detects and associates hybrid genomes to appropriate the groups. Additional putative hybrids include *R. humulifolius,* with possible parentage from species in subg. *Idaeobatus* and *Cylactis*, and *R. macraei,* with putative progenitors from *Idaeobatus* and a species, such as *R. ursinus*, from subg. *Rubus* (40). Long read sequence data and the assembly of haplotypes would give additional insight into difficult to classify polyploid, hybrid species like *R. macraei* and *R. chamaemorus* (59, 60). Haplotype sequencing could allow direct analysis of the evolutionary history of different subgenomes in these putative hybrid species. Each subgenome would be treated as a separate branch on the phylogeny. Instead of hybrids showing an intermediate relationship with progenitors, as in our analysis, subgenome sequences would group directly with parental species. However, our use of hundreds of loci, multiple analysis methods, and assessment of phylogenetic signal supporting internal nodes enabled a critical assessment of the broad evolutionary history of *Rubus*.

Our molecular analysis and dating approach estimated the biogeographical patterns in *Rubus.* The most recent common ancestor was likely distributed in North America. During the early Miocene, lineages likely migrated from North America to Asia and Europe over the Bering land bridge. Migrations to South America occurred during the formation of the Panamanian Isthmus in the mid- to late Miocene, and long-distance dispersal events may have allowed *Rubus* to spread from South America to Australia and New Zealand. During the middle and late Miocene the genus diversified greatly in Asia, Europe, South America and Oceania. Whole genome duplication events occurred producing higher ploidy species on multiple continents. Cooling temperatures and glaciation isolated Central American populations from North America, and may have created conditions beneficial to the formation of apomictic polyploids in Europe.

While our research sets the stage for reassessing *Rubus* subtaxa, i.e., subgenera or sections, a thorough morphological evaluation of multiple accessions of species across the genus must follow to identify useful synapomorphies for taxonomic redefinition.

## Methods

### Samples and ethics statement

Samples designated by a PI number (Table 1), were obtained from the USDA ARS NCGR according to rules of the International Treaty on Plant Genetic Resources for Food and Agriculture (61). DNA from leaf samples without PI numbers were obtained by LA through field work, and exchange from international botanical gardens and herbaria (Table 1).

### Sampling and DNA extraction

We sampled 94 accessions, representing 87 wild *Rubus*, three cultivars (*R.* hybrid ‘Logan’, ‘Boysen’, ‘Marion’), and outgroup *Waldsteinia fragarioides* (Table 1). *Rubus* is sister to the clade containing *Waldsteinia* in the phylogeny of Rosaceae estimated by Potter et al. (62) and Xiang et al. (25). Twenty-six species are from subg. *Idaeobatus*, 24 are from subg. *Rubus* and other subgenera are represented by 1-9 species each (Additional file S5). ‘Logan’, ‘Boysen’, and ‘Marion’ were sampled because they are economically important hybrid cultivars with known percentages of blackberry and raspberry parentage. ‘Logan’ is comprised of 50% blackberry/50% raspberry species; ‘Boysen’, an offspring of ‘Logan’, is 75% blackberry/25% raspberry; and ‘Marion’ is 69% blackberry/31% raspberry (3, 4). Genomic DNA was isolated from fresh leaves frozen at −80 °C, leaves dried on silica gel desiccant, or herbarium specimens (17, 63–65) using a modified CTAB (hexadecyltrimethylammonium bromide) extraction method (66).

### Target enrichment probe design

Targets were developed from the *Rubus occidentalis* genome v1 assembly (67) and a conserved set of loci from *Fragaria vesca*, *Malus* × *domestica* and *Prunus persica* (68). Exon sequences were extracted from the *R. occidentalis* transcriptome assembly (67). Only those exons ≥ 80 bp, with GC content between 30 and 70%, and with one BLAST hit to the *R. occidentalis* genome over 50% of the exon length and with ≥ 90% identity were used for bait development. In total, probes were synthesized by MYcroarray (now Arbor Biosciences, Ann Arbor, MI, USA) for 8,963 exons from 926 genes. Due to a bioinformatics error, the *R. occidentalis* exon sequences from which probes were created were cropped into 60 bp sequences separated by 20 bp gaps before submission to MYcroarray. The 120-mer baits synthesized by MYcroarray with 1x tiling corresponded to 140 bp of genome sequence. Despite this, hybridization with the *R. occidentalis-*derived probes was successful for nearly all study samples.

Conserved loci from *F. vesca*, *Malus × domestica* and *P. persica* genomes were selected for their usefulness in comparative genomic studies across Rosaceae as describedListon (68). Briefly, single copy loci shared between the *F. vesca* and *P. persica* genomes were identified. The corresponding genes were extracted from the *Malus × domestica* genome, where there were often two gene copies due to the allopolyploid ancestry of the domesticated apple. The gene sequence with the fewest ambiguous bases or polymorphic sites was selected. Genes were filtered based on their phylogenetic utility (≥ 960 bp, > 85% pairwise sequence similarity between the three genomes) and to maximize the success of target capture (exons ≥ 80 bp, GC content > 30% or < 70%, < 90% sequence similarity to other target exons in the same genome). This resulted in 257 genes; probes were designed for the copies of these genes originating from *F. vesca*.

### Library preparation

Genomic DNA was quantified with PicoGreen (ThermoFisher Scientific, Waltham, MA, USA) and quality checked using agarose gel electrophoresis. To prepare for library construction, 400 ng of input DNA was sonicated for 5-10 minutes using a Diagenode BioRuptor Sonicator (Denville, NJ, USA). After an initial 5 minutes of sonication, samples were sized using gel electrophoresis and sonicated an additional 1-5 minutes as necessary to achieve the desired 200 bp average insert size. If DNA bands were very faint after the first round of sonication, a new aliquot of the sample with 600-800 ng of input DNA was prepared and sonicated. Sonicated samples were cleaned using Qiaquik PCR purification columns (QIAGEN, Valencia, CA, USA) to eliminate low molecular weight fragments. Genomic libraries were prepared using the NEBNext Ultra DNA Library Prep Kit with NEBNext Multiplex Oligos for Illumina (New England Biolabs, Ipswich, MA, USA) to enable multiplexed sequencing. Size selection for 200 bp fragments was done after adaptor ligation using AMPure (Agencourt Bioscience Corporation, A Beckman Coulter Company, Beverly, MA, USA) beads at a 0.55:1 ratio with the sample. Libraries were amplified for 8 PCR cycles and cleaned with AMPure beads at a 1:1 ratio with the sample before being quantified with PicoGreen. A subset of libraries was quality checked with the Agilent Bioanalyzer (Agilent Technologies, Santa Clara, CA, USA) at Oregon State University’s (OSU) Center for Genome Research and Biocomputing (CGRB).

To prepare for in-solution hybridization, samples were divided into four pools of 24 samples containing 20 ng of each library. MYcroarray MYbaits (Arbor Biosciences, Ann Arbor, USA) protocol version 1.3.8 was followed for sequence enrichment. The resulting pools were quantified using Qubit and qPCR, pooled again in equimolar amounts and sequenced with 100 bp paired end reads in one Illumina® HiSeq ^TM^ 2000 lane at the CGRB. Libraries were demultiplexed using the Illumina pipeline.

### Sequence assembly

Bases with a quality score under Q20 were trimmed from the right and left side of reads with BBduk; reads shorter than 25 bp after trimming were discarded (69). Loci were assembled with HybPiper v. 1.2 using sequence read files and a target sequence reference file from which probes were designed (45). To replace the missing 20 bp sequences from the *Rubus* baits in this target reference file, the 60 bp target fragments used in probe synthesis were first mapped against the *R. occidentalis* genome with BBMap. Then, Bedtools v. 2.25.0 was used to extract contiguous sequences for each exon (69, 70). Exons for each gene were then concatenated to create the final target sequence reference. HybPiper creates bins based on reads by target sequence using BWA (71). The reads are then assembled with SPAdes into contigs using the target sequence as a reference (72, 73). Output sequences were either assembled exons or supercontigs, which could include noncoding sequences such as introns, 5’ UTR, and 3’ UTR sequences obtained from genomic libraries during hybridization.

Exons and supercontig sequences were each aligned with MAFFT v. 7.402. Alignment sites with gaps in more than 20% of sequences were removed with TrimAl v. 1.2rev59 (74).

### Phylogenetic analyses of nuclear loci

The maximum likelihood phylogeny was estimated twice for each locus, once with the exon sequences and secondly with the supercontig sequence data. We used RAxML v. 8.1.21 to conduct a bootstrap search with up to 1000 replicates (-#autoMRE or -#1000 option) and estimate the maximum likelihood phylogeny for each gene [option –f a; Stamatakis (75)]. The best fit model of evolution (GTRGAMMA or GTRGAMMAI) was determined with PartitionFinder v. 2.1.1 for the exon sequences of each gene. This same model was also used for supercontig sequence analyses (76). Phylogenies were estimated for two sets of taxa: one containing only diploids and the other containing all taxa polyploids and diploids. Thus, for each gene, a phylogeny was estimated for the following datasets: diploid exons, diploid supercontig sequences, all taxa exons, and all taxa supercontig sequences.

To prevent ambiguous placement of taxa in a tree resulting from insufficient phylogenetic signal, RogueNaRok v. 1.0 was used with default settings to identify such “rogue” taxa for each locus using bootstrapped RAxML trees (77). Rogue taxa were eliminated from sequence alignments and gene trees were re-estimated with RAxML.

Species phylogenies were estimated under the multi-species coalescent model using ASTRAL-II v. 4.10.12 and SVDQuartets implemented in PAUP* 4.0 (78–80). ASTRAL-II and SVDQuartets both use relationships between quartets of taxa to estimate the overall species tree. ASTRAL-II identified the species tree that shares the maximum number of quartet trees with the 941 gene trees estimated with RAxML (78). Local posterior probability support values were calculated as these have been shown to be highly precise compared with multi-locus bootstrapping (81). SVDQuartets randomly sampled 100,000 possible quartets of taxa and used SNPs from the concatenated sequence alignments to score each possible split in the quartets [100 boostrap replicates; (78)]. The best scoring splits were assembled into a species phylogeny in PAUP* using QFM (80, 82).

Branch support for phylogenies with the highest likelihood for each concatenated sequence alignment were also evaluated using Quartet Sampling (83). This method evaluates the topological relationship between quartets of taxa using an input phylogeny and a molecular alignment partitioned by gene. Unlike bootstrap values, this method can distinguish if the data supporting internal branches is strongly discordant or lacking signal (83). Quartet Sampling produces three main scores, quartet concordance (QC), quartet differential (QD), and quartet informativeness (QI) for each node. Quartet concordance describes how often concordant quartets, which show the same splits and sister relationships between clades, are inferred. Scores ≥ 0.5 indicate strong support for the concordant topology. Quartet differential measures how often quartets with discordant topologies are inferred. This measure can indicate if a dataset shows strong evidence for an alternate evolutionary history at a node. Scores ∼1 indicate that no alternate topology is strongly favored. Quartet informativeness measures the proportion of replicates that are informative for a node. Scores =1 indicate that all replicates were informative while scores =0 indicate that none were informative.

### Network analysis

Because we anticipated high levels of ILS and hybridization in this dataset, we estimated unrooted super networks to visualize incongruences among exon or supercontig sequence gene trees and identify putative hybrid taxa using SuperQ v. 1.1 with the Gurobi optimizer and a balanced linear secondary objective function (84). In this method, input gene trees (identical to gene trees used in ASTRAL-II analyses) are broken down into quartets and reassembled into a network where edge lengths indicate the frequency of each split in the gene tree set.

### Dating for Phylogenetic estimation

ASTRAL-II-generated topologies from genes estimated using exon sequences were used for dating. Branch lengths per site substitution rates were estimated over the ASTRAL-II topology for all taxa using RAxML [-f e option, GTRGAMMA model of evolution; Stamatakis (75)] and the corresponding concatenated alignment of exon sequences. Phylogenies were dated with r8s version 1.80 using the penalized likelihood method and the truncated Newton algorithm with a smoothing parameter estimated using cross validation (85, 86). The age of the root node was constrained to 56.93-65.66 Ma based on the age of this node estimated from plastid sequences (26).

### Biogeographic analyses

Data were collected for the continent of origin for each sample (Table 1). Ancestral ranges were estimated with BioGeoBEARS version 1.1 over ultrametric dated phylogenies resulting from r8s using Dispersal-Extinction-Cladogenesis (DEC) and DEC+*j* likelihood models (87, 88).The parameter *j* incorporates founder-event speciation or long distance dispersal events (87, 89).

### Chloroplast sequence extraction and analysis

Reads for each sample were mapped to the *R. occidentalis* chloroplast reference genome (67) edited with BBMap to contain only one copy of the inverted repeat (67, 69). Consensus chloroplast sequences from a reduced read set of up to 100,000 mapped reads were extracted using Geneious v. 9.1.7 with Ns inserted at sites with no sequence coverage (90). Consensus sequences were aligned with MAFFT using auto settings (91). Alignment sites with missing data in over 20% of samples were stripped using Geneious v. 9.1.7 (90). The maximum likelihood phylogeny was estimated with RAxML using up to 1000 bootstrap replicates [-#autoMRE and –f a options; Stamatakis (75)] under the GTRGAMMAI model of evolution. Rogue taxa were identified with RogueNaRok and removed from the alignment (77). RAxML was subsequently run to estimate the final maximum likelihood phylogeny.

## List of abbreviations

CGRB: Center for Genome Research and Biocomputing
DEC: Dispersion extinction cladogram
ILS: Incomplete lineage sorting
Ma: Million years ago
MRCA: Most recent common ancestor
OSU: Oregon State University
PI: Plant Information number for the USDA
QC: quartet concordance
QD: quartet differential
QI: quartet informativeness
USDA: United States Department of Agriculture

## Data accessibility

Sequence alignments and phylogenies are available at (OSU scholars archive link).

OSU Scholars Archive

Short Read Accessions (SRA): PRJNA510412

Temporary Submission ID: SUB4927845

Release date: 2020-01-17

## Additional files

Additional file 1 Table S1. Success of target capture for each *Rubus* sample. Statistics except average read depth calculated with HybPiper script hybpiper_stats.py. Total number of target genes is 1173. Samples marked with an asterisk did not sequence well and were not included in phylogenetic analyses.

Additional file 2 Table S2. Alignment statistics for analyses for exon and supercontigs (S, exon + noncoding sequences) for diploid and all taxa *Rubus* datasets. Samples marked with an asterisk did not sequence well and were not included in phylogenetic analyses. E = exon sequence alignments. Ungapped lengths are reported.

Additional file 3 Table S3. Chloroplast sequence statistics. *Rubus* samples marked with an asterisk either failed sequencing or were removed from the analysis as rogue taxa by RogueNaRok. Average read depth was calculated from mapping up to 100,000 reads against the *R. occidentalis* reference chloroplast sequence edited to include one copy of the inverted repeat region.

Additional file 4 Table S4. Putative *Rubus* hybrid groups and species. Hybrid origins supported by discordance in nuclear phylogenies and networks. Maternal progenitors inferred from the chloroplast phylogeny.

Additional file 5 Table S5. Number of studied accessions classified in each subgenus. “Other hybrids” include two *R. ursinus* × *armeniacus* hybrids.

## Declarations

The authors declare that they have ethics approval and consent to participate; that they have consent for publication; that the data and material are available; that they have no competing interests.

## Authors’ contributions

KC contributed to the laboratory work, data analyses, and manuscript writing; LA, BS, and JB contributed to the laboratory work and manuscript review; TM and DB contributed to data analysis and manuscript review; AL, LA, NB, and KH conceived the study and contributed to the analysis and manuscript preparation; all authors have read and approved the final manuscript.

## Acknowledgements

We appreciate the technical assistance of the Center for Genome Research and Biocomputing at Oregon State University for Illumina® sequencing. We thank M. Dossett, R. Cronn, K. Weitemier, R. Schmickl, J.C. Lee, R. Meiers, C. Mulch, A.M. Nyberg, M. Peterson, M. Clark, K.J. Vining, M.L. Worthington, M.H. Yin, J.D. Zurn, J.R. Clark, C.E. Finn for technical assistance and meaningful discussion on this manuscript. This work was supported by grants from USDA ARS CRIS 2072-21000-044-00D and 2072-21000-049-00D and NSF KY EPSCoR National Laboratory Initiative 019-14 and NSF DEB 0236166 grants awarded to LAA for support of this research.

## References

1. Hytönen T, Graham J, Harrison R. The Genomes of Rosaceous Berries and Their Wild Relatives. Springer; 2018.

2. FAO. Food and Agriculture Organization of the United Nations, Crop Statistical Databases 2019 [Accessed 13 February 2019]. Available from: http://www.fao.org/faostat/en/#data/QC.

3. Jennings DL. Raspberries and blackberries: their breeding, diseases and growth. London, UK: Academic Press; 1988.

4. Thompson MM. Survey of chromosome numbers in *Rubus* (Rosaceae: Rosoideae). Annals of the Missouri Botanical Garden. 1997:128–64.

5. Thompson MM. Chromosome numbers of *Rubus* species at the National Clonal Germplasm Repository. HortScience. 1995;30(7):1447–52.

6. Sochor M, Vašut RJ, Sharbel TF, Trávníček B. How just a few makes a lot: speciation via reticulation and apomixis on example of European brambles (Rubus subgen. Rubus, Rosaceae). Molecular phylogenetics and evolution. 2015;89:13–27.

7. Alice LA, Eriksson T, Eriksen B, Campbell CS. Hybridization and Gene Flow between Distantly Related Species of *Rubus* (Rosaceae): Evidence from Nuclear Ribosomal DNA Internal Transcribed Spacer Region Sequences. Systematic Botany. 2001;26(4):769–78.

8. Wang Y, Wang X, Chen Q, Zhang L, Tang H, Luo Y, et al. Phylogenetic insight into subgenera Idaeobatus and Malachobatus (Rubus, Rosaceae) inferring from ISH analysis. Molecular cytogenetics. 2015;8(1):11.

9. Focke WO. Species ruborum. Monographiae generis rubi prodromus. Stuttgart: Stuttgart, E. Schweizerbart; 1910.

10. Focke WO. Species ruborum. Monographiae generis rubi prodromus. Stuttgart: Stuttgart, E. Schweizerbart; 1911.

11. Focke WO. Species ruborum. Monographiae generis rubi prodromus. Stuttgart: Stuttgart, E. Schweizerbart; 1914.

12. Lu L-T, Bufford D. eFlora China Rubus. Flora of China. 2003;9(Rosaceae).

13. Bean AR. Revision of *Rubus* subgenus *Micranthobatus* (Fritsch) Kalkman (Rosaceae) in Australia. Austrobaileya. 1995;4(3):321–8.

14. Bean AR. A revision of *Rubus* subg. *Malachobatus* (Focke) Focke and *Rubus* subg. *Diemenicus* AR Bean (Rosaceae) in Australia. Austrobaileya. 1997:39–51.

15. Bailey LH. Species batorum: the genus *Rubus* in North America: Bailey Hortorium of the The New York State College of Agriculture at Cornell University; 1941.

16. Lu L, Boufford D, E. Rubus L. Flora of China. 2003;9:195–286.

17. Alice LA, Campbell CS. Phylogeny of *Rubus* (Rosaceae) Based on Nuclear Ribosomal DNA Internal Transcribed Spacer Region Sequences. American Journal of Botany. 1999;86(1):81–97.

18. Wang Y, Chen Q, Chen T, Tang H, Liu L, Wang X. Phylogenetic Insights into Chinese *Rubus* (Rosaceae) from Multiple Chloroplast and Nuclear DNAs. Frontiers in Plant Science. 2016;7(968).

19. Mimura M, Mishima M, Lascoux M, Yahara T. Range shift and introgression of the rear and leading populations in two ecologically distinct Rubus species. BMC evolutionary biology. 2014;14(1):209.

20. Bammi RK, Olmo HP. Cytogenetics of *Rubus*. V. Natural hybridization between *R. procerus* PJ Muell. and *R. laciniatus* Willd. Evolution. 1966;20(4):617–33.

21. Rabeler RK, Hartman RL. Flora of North America. From Internet. 2007.

22. Yang JY, Pak J-H. Phylogeny of Korean *Rubus* (Rosaceae) based on ITS (nrDNA) and trnL/F intergenic region (cpDNA). Journal of Plant Biology. 2006;49(1):44–54.

23. Maddison WP. Gene trees in species trees. Systematic biology. 1997;46(3):523–36.

24. Lu L-T. Study on the Genus Rubus of China. Chih wu fen lei hsueh pao= Acta Phytotaxonomica Sinica. 1983.

25. Xiang Y, Huang C-H, Hu Y, Wen J, Li S, Yi TS, et al. Evolution of Rosaceae fruit types based on nuclear phylogeny in the context of geological times and genome duplication. Molecular biology and evolution. 2016;34(2):262–81.

26. Zhang SD, Jin JJ, Chen SY, Chase MW, Soltis DE, Li HT, et al. Diversification of Rosaceae since the Late Cretaceous based on plastid phylogenomics. New Phytologist. 2017;214(3):1355–67.

27. Graham A. The role of land bridges, ancient environments, and migrations in the assembly of the North American flora. Journal of Systematics and Evolution. 2018.

28. Tiffney BH. Perspectives on the origin of the floristic similarity between eastern Asia and eastern North America. Journal of the Arnold Arboretum. 1985;66(1):73–94.

29. Milne RI. Northern hemisphere plant disjunctions: a window on Tertiary land bridges and climate change? Annals of Botany. 2006;98(3):465–72.

30. Wen J. Evolution of Eastern Asian and Eastern North American Disjunct Distributions in Flowering Plants. Annual Review of Ecology and Systematics. 1999;30(1):421–55.

31. Wen J. Evolution of Eastern Asian and Eastern North American biogeographic disjunctions: a few additional issues. International Journal of Plant Sciences. 2001;162(S6):S117–S22.

32. Wen J, Nie Z-L, Ickert-Bond SM. Intercontinental disjunctions between Eastern Asia and Western North America in vascular plants highlight the biogeographic importance of the Bering land bridge from late Cretaceous to Neogene. Journal of Systematics and Evolution. 2016;54(5):469–90.

33. O’Dea A, Lessios HA, Coates AG, Eytan RI, Restrepo-Moreno SA, Cione AL, et al. Formation of the Isthmus of Panama. Science Advances. 2016;2(8):e1600883.

34. Weitemier K, Straub SC, Cronn RC, Fishbein M, Schmickl R, McDonnell A, et al. Hyb-Seq: Combining target enrichment and genome skimming for plant phylogenomics. Applications in Plant Sciences. 2014;2(9):1400042.

35. Folk RA, Mandel JR,. Freudenstein JV. Ancestral gene flow and parallel organellar genome capture result in extreme phylogenomic discord in a lineage of angiosperms. Systematic biology. 2017;66(3):320–37.

36. Morales-Briones DF, Liston A, Tank DC. Phylogenomic analyses reveal a deep history of hybridization and polyploidy in the Neotropical genus Lachemilla (Rosaceae). New Phytologist. 2018;218(4):1668–84.

37. Dillenberger MS, Wei N, Tennessen JA, Ashman TL, Liston A. Plastid genomes reveal recurrent formation of allopolyploid Fragaria. American Journal of Botany. 2018.

38. Carter KA. Phylogenetic estimation and ancestral state reconstruction of Rubus (Rosaceae) using target capture sequencing Corvallis, Oregon: Oregon State University; 2018.

39. Chou J, Gupta A, Yaduvanshi S, Davidson R, Nute M, Mirarab S, et al. A comparative study of SVDquartets and other coalescent-based species tree estimation methods. BMC genomics. 2015;16(10):S2.

40. Morden CW, Gardner DE, Weniger DA. Phylogeny and biogeography of pacific Rubus subgenus Idaeobatus (Rosaceae) species: Investigating the origin of the endemic Hawaiian raspberry R. macraei. Pacific Science. 2003;57(2):181–97.

41. Gleason H, Cronquist A. Manual of vascular plants of northeastern North America and adjacent Canada. New York Botanical Garden, Bronx, New York, USA. 1991.

42. Hummer KE, Alice LA. Small Genomes in Tetraploid *Rubus* L.(Rosaceae) from New Zealand and Southern South America. Journal of the American Pomological Society. 2017;71(1):2–7.

43. Pankhurst R. Rosaceae. ∼En:∼ Stevens, W.D., C. Ulloa, A. Pool & O.M. Montiel (eds.). Flora de Nicaragua. Monographs in Systematic Botany from the Missouri Botanical Garden. 2001;85(3):2202–6.

44. Standley PC, Steyermark JA. Flora of Guatemala. Flora of Guatemala. 1946.

45. Johnson MG, Gardner EM, Yang L, Medina R, Goffinet B, Shaw AJ, et al. HybPiper: Extracting coding sequence and introns for phylogenetics from high-throughput sequencing reads using target enrichment. Applications in Plant Sciences. 2016;4(7):apps.1600016.

46. Seehausen O. Hybridization and adaptive radiation. Trends in ecology & evolution. 2004;19(4):198–207.

47. Hall HK, Funt RC. Blackberries and their Hybrids. Crop Production Science in Horticulture: CABI; 2017.

48. Michael K. Clarification of basal relationships in *Rubus* (Rosaceae) and the origin of *Rubus chamaemorus*. Bowling Green, Kentucky: Westren Kentucky University; 2006.

49. Martinussen I, Uleberg E, Sønsteby A, Sønstebø JH, Graham J, Vivian-Smith A. Genomic survey sequences and the structure of the Rubus chamaemorus L. genome as determined by ddRAD tags 2013.

50. Howarth DG, Gardner DE, Morden CW. Phylogeny of *Rubus* Subgenus *Idaeobatus* (Rosaceae) and its Implications Toward Colonization of the Hawaiian Islands. Systematic Botany. 1997;22(3):433–41.

51. Kalkman C. The phylogeny of the Rosaceae. Botanical Journal of the Linnean Society. 1988;98(1):37–59.

52. Nie Z-L, Sun H, Manchester SR, Meng Y, Luke Q, Wen J. Evolution of the intercontinental disjunctions in six continents in the Ampelopsis clade of the grape family (Vitaceae). BMC evolutionary biology. 2012;12:17-.

53. Wang Y. Relationships among Rubus (Rosaceae) species used in traditional Chinese medicine. 2011.

54. Williamson PG. Selection or constraint? A proposal on the mechanism for stasis. Rates of evolution Allen and Unwin, London. 1987:129–42.

55. Parks CR, Wendel JF. Molecular divergence between Asian and North American species of Liriodendron (Magnoliaceae) with implications for interpretation of fossil floras. American Journal of Botany. 1990;77(10):1243–56.

56. Milne RI, Abbott RJ. The origin and evolution of Tertiary relict floras. Advances in Botanical Research. 2002;38:281–314.

57. Graham A. History of the vegetation: cretaceous (Maastrichtian)-Tertiary. Flora of North America. 1993;1:57–70.

58. Lawver LA, Gahagan LM. Evolution of Cenozoic seaways in the circum-Antarctic region. Palaeogeography, Palaeoclimatology, Palaeoecology. 2003;198(1):11–37.

59. Kamneva OK, Syring J, Liston A, Rosenberg NA. Evaluating allopolyploid origins in strawberries (Fragaria) using haplotypes generated from target capture sequencing. BMC evolutionary biology. 2017;17(1):180.

60. Dauphin B, Grant JR, Farrar DR, Rothfels CJ. Rapid allopolyploid radiation of moonwort ferns (Botrychium; Ophioglossaceae) revealed by PacBio sequencing of homologous and homeologous nuclear regions. Molecular phylogenetics and evolution. 2018;120:342–53.

61. ITPGR. International Treaty on Plat Genetic Resources for Food and Agriculture: Food and Agriculture Organization of the United Nations; 2019 [Available from: http://www.fao.org/plant-treaty/en/.

62. Potter D, Eriksson T, Evans R, Oh S-H, Smedmark JEE, Morgan RD, et al. Phylogeny and classification of Rosaceae2007. 5–43 p.

63. Holmgren PK, Holmgren NH, Barnett LC. Index Herbariorum, Part I: The herbaria of the world,. New York, New York: New York Botanical Garden; 1990.

64. Alice L, Dodson T, Sutherland B. Diversity and relationships of Bhutanese Rubus. Acta horticulturae. 2008.

65. Morden CW, Caraway V, Motley TJ. Development of a DNA library for native Hawaiian plants. 1996.

66. Doyle J, Doyle JL. Genomic plant DNA preparation from fresh tissue-CTAB method. Phytochem Bull. 1987;19(11):11–5.

67. VanBuren R, Bryant D, Bushakra JM, Vining KJ, Edger PP, Rowley ER, et al. The genome of black raspberry (*Rubus occidentalis*). The Plant Journal. 2016;87(6):535–47.

68. Liston A. 257 nuclear genes for Rosaceae phylogenomics2014.

69. Bushnell B. BBTools software package. URL http://sourceforgenet/projects/bbmap. 2014.

70. Quinlan AR. BEDTools: the Swiss-army tool for genome feature analysis. Current protocols in bioinformatics. 2014;47(1):11.2. 1-.2. 34.

71. Li HT, Durbin R. Fast and accurate short read alignment with Burrows–Wheeler transform. Bioinformatics. 2009;25(14):1754–60.

72. Li H. Aligning sequence reads, clone sequences and assembly contigs with BWA-MEM. arXiv preprint arXiv:13033997. 2013.

73. Bankevich A, Nurk S, Antipov D, Gurevich AA, Dvorkin M, Kulikov AS, et al. SPAdes: a new genome assembly algorithm and its applications to single-cell sequencing. Journal of computational biology. 2012;19(5):455–77.

74. Capella-Gutiérrez S, Silla-Martínez JM, Gabaldón T. trimAl: a tool for automated alignment trimming in large-scale phylogenetic analyses. Bioinformatics. 2009;25(15):1972–3.

75. Stamatakis A. RAxML version 8: a tool for phylogenetic analysis and post-analysis of large phylogenies. Bioinformatics. 2014;30(9):1312–3.

76. Lanfear R, Calcott B, Ho SY, Guindon S. PartitionFinder: combined selection of partitioning schemes and substitution models for phylogenetic analyses. Molecular biology and evolution. 2012;29(6):1695–701.

77. Aberer AJ, Krompass D, Stamatakis A. Pruning Rogue Taxa Improves Phylogenetic Accuracy: An Efficient Algorithm and Webservice. Systematic Biology. 2013;62(1):162–6.

78. Mirarab S, Warnow T. ASTRAL-II: coalescent-based species tree estimation with many hundreds of taxa and thousands of genes. Bioinformatics. 2015;31(12):i44–i52.

79. Chifman J, Kubatko L. Quartet Inference from SNP Data Under the Coalescent Model. Bioinformatics. 2014;30(23):3317–24.

80. Swofford D. PAUP* ver 4.0. b10. Phylogenetic Analysis Using Parsimony and Other Methods Sunderland, MA: Sinauer Associates, Sunderland. 2003.

81. Sayyari E, Mirarab S. Fast Coalescent-Based Computation of Local Branch Support from Quartet Frequencies. Molecular Biology and Evolution. 2016;33(7):1654–68.

82. Reaz R, Bayzid MS, Rahman MS. Accurate phylogenetic tree reconstruction from quartets: A heuristic approach. PLoS One. 2014;9(8):e104008.

83. Pease JB, Brown JW, Walker JF, Hinchliff CE, Smith SA. Quartet Sampling distinguishes lack of support from conflicting support in the green plant tree of life. American journal of botany. 2018;105(3):385–403.

84. Grunewald S, Spillner A, Bastkowski S, Bogershausen A, Moulton V. SuperQ: computing supernetworks from quartets. IEEE/ACM Transactions on Computational Biology and Bioinformatics (TCBB). 2013;10(1):151–60.

85. Sanderson MJ. Estimating absolute rates of molecular evolution and divergence times: a penalized likelihood approach. Molecular biology and evolution. 2002;19(1):101–9.

86. Sanderson MJ. r8s: inferring absolute rates of molecular evolution and divergence times in the absence of a molecular clock. Bioinformatics. 2003;19(2):301–2.

87. Matzke NJ. Model selection in historical biogeography reveals that founder-event speciation is a crucial process in island clades. Systematic Biology. 2014;63(6):951–70.

88. Ree RH, Smith SA. Maximum likelihood inference of geographic range evolution by dispersal, local extinction, and cladogenesis. Systematic biology. 2008;57(1):4–14.

89. Matzke NJ. BioGeoBEARS: biogeography with Bayesian (and likelihood) evolutionary analysis in R scripts. R package, version 11. 2013;1:2013.

90. Kearse M, Moir R, Wilson A, Stones-Havas S, Cheung M, Sturrock S, et al. Geneious Basic: an integrated and extendable desktop software platform for the organization and analysis of sequence data. Bioinformatics. 2012;28(12):1647–9.

91. Katoh K, Standley DM. MAFFT Multiple Sequence Alignment Software Version 7: Improvements in Performance and Usability. Molecular Biology and Evolution. 2013;30(4):772–80.

92. Sutherland B. Phylogenetics of Rubus ursinus and R. macraai (Rosaceae): Evidence of hybrid origin. Student Honors Theses. 2005:186.

93. Barneby RC. Flora of bhutan, including a record of plants from sikkim. Vol. 1, Part 3. Brittonia. 1988;40(3):289-.

94. Romoleroux K, Nllgaard B, Harling G, Andersoon L. Flora of Ecuador: 79. Rosaceae; 81. Connaraceae: Department of Systematic Botany; 1996.

95. Meng R, Finn CE. Determining ploidy level and nuclear DNA content in Rubus by flow cytometry. Journal of the American society for Horticultural Science. 2002;127(5):767–75.

96. Hummer KE, Bassil NV, Alice LA, editors. *Rubus* ploidy assessment. XI International Rubus and Ribes Symposium 1133; 2015.

