## Supplemental Files for "Target Capture Sequencing Unravels *Rubus* Evolution"

Additional Files: Tables supplemental

Additional File 1: Table S1. Success of target capture for each *Rubus* sample. Statistics except average read depth calculated with HybPiper script hybpiper_stats.py. Total number of target genes is 1173. Samples marked with an asterisk did not sequence well and were not included in phylogenetic analyses.

| **­** | **Number of reads (Mbp)** | **Reads mapped to targets (Millions)** | **% Reads on target** | **Number of target genes with sequences** | **Number of target sequences ≥ 75% of target length** | **Average read depth over all targets** |
| --- | --- | --- | --- | --- | --- | --- |
| **Average** | **2.7** | **1.9** | **72%** | **1090** | **988** | **66.8** |
| Boysen | 3.0 | 2.0 | 67% | 1140 | 1054 | 71.1 |
| Logan | 5.3 | 3.9 | 74% | 1149 | 1119 | 137.9 |
| Marion | 2.6 | 1.9 | 74% | 1145 | 1087 | 66.2 |
| *R. acanthophyllus* | 4.0 | 3.3 | 81% | 1148 | 1103 | 114.2 |
| *R. alexeterius* | 1.4 | 1.2 | 87% | 1140 | 1065 | 42.6 |
| *R. allegheniensis* | 0.2 | 0.2 | 72% | 778 | 66 | 6.0 |
| *R. amphidasys* | 4.9 | 3.1 | 63% | 1146 | 1094 | 110.0 |
| *R. arcticus* | 3.0 | 2.4 | 80% | 1146 | 1087 | 83.3 |
| *R. argutus* | 3.3 | 2.4 | 73% | 1141 | 1083 | 85.6 |
| *R. armeniacus* | 1.3 | 1.0 | 74% | 1110 | 929 | 33.8 |
| *R. assamensis* | 7.1 | 5.5 | 77% | 1143 | 1104 | 192.0 |
| *R. australis* | 7.0 | 5.5 | 79% | 1148 | 1097 | 194.6 |
| *R. bifrons* | 2.0 | 1.5 | 75% | 1133 | 1039 | 53.5 |
| *R. caesius* | 5.6 | 3.5 | 62% | 1148 | 1102 | 122.2 |
| *R. calophyllus* | 2.1 | 1.6 | 79% | 1137 | 1033 | 57.7 |
| *R. calycinus* | 1.5 | 1.2 | 78% | 1134 | 994 | 41.6 |
| *R. canadensis* | 4.9 | 3.6 | 73% | 1146 | 1092 | 125.8 |
| *R. caucasicus* | 1.1 | 0.9 | 84% | 1126 | 999 | 32.5 |
| *R. chamaemorus* | 0.6 | 0.5 | 84% | 1081 | 639 | 17.3 |
| *R. coreanus* | 4.7 | 3.8 | 80% | 1152 | 1112 | 132.2 |
| *R. coriifolius* | 1.2 | 0.9 | 70% | 1101 | 902 | 30.5 |
| *R. crataegifolius* | 4.2 | 2.7 | 64% | 1143 | 1077 | 93.9 |
| *R. cuneifolius* | 2.3 | 1.8 | 77% | 1138 | 1055 | 61.5 |
| *R. deliciosus* | 2.0 | 1.5 | 74% | 1139 | 1064 | 52.1 |
| *R. ellipticus* | 2.2 | 1.7 | 75% | 1141 | 1070 | 58.1 |
| *R. eriocarpus* | 3.1 | 2.5 | 80% | 1148 | 1098 | 88.2 |
| *R. flagellaris* | 5.9 | 4.8 | 81% | 1149 | 1121 | 167.1 |
| *R. fockeanus* | 5.4 | 3.8 | 70% | 1149 | 1108 | 132.0 |
| *R. geoides* | 5.4 | 4.1 | 77% | 1152 | 1111 | 144.5 |
| *R. glabratus* | 5.1 | 3.4 | 67% | 1147 | 1106 | 121.1 |
| *R. glaucus* | 1.4 | 1.1 | 77% | 1119 | 986 | 36.9 |
| *R. gunnianus* | 2.9 | 2.1 | 74% | 1140 | 1074 | 75.4 |
| *R. hawaiensis* | 1.3 | 1.0 | 81% | 1127 | 1009 | 35.6 |
| *R. hispidus** | 0.0 | 0.0 | 82% | 3 | - | 0.4 |
| *R. humulifolius* | 1.0 | 0.8 | 83% | 1119 | 978 | 28.2 |
| *R. ichangensis* | 6.1 | 4.9 | 80% | 1146 | 1109 | 170.4 |
| *R. idaeus* | 2.9 | 2.0 | 69% | 1151 | 1099 | 71.5 |
| *R. illecebrosus* | 4.9 | 3.3 | 67% | 1143 | 1091 | 114.8 |
| *R. innominatus* | 2.0 | 1.6 | 79% | 1135 | 1063 | 54.5 |
| *R. irenaeus* | 5.2 | 3.9 | 76% | 1149 | 1103 | 138.4 |
| *R. laciniatus* | 1.9 | 1.4 | 76% | 1132 | 1038 | 50.7 |
| *R. laegardii* | 1.2 | 1.1 | 87% | 1131 | 994 | 37.0 |
| *R. lambertianus* | 2.5 | 2.0 | 79% | 1137 | 1066 | 69.2 |
| *R. lasiococcus* | 0.8 | 0.6 | 78% | 1107 | 902 | 22.2 |
| *R. lasiostylus* | 4.9 | 3.3 | 69% | 1148 | 1101 | 117.5 |
| *R. leucodermis* | 3.7 | 2.4 | 65% | 1154 | 1111 | 84.3 |
| *R. lineatus* | 2.3 | 1.8 | 80% | 1140 | 1054 | 64.3 |
| *R. loxensis* | 1.4 | 1.0 | 76% | 1112 | 946 | 36.5 |
| *R. macraei* | 3.1 | 2.5 | 81% | 1146 | 1095 | 87.3 |
| *R. moorei* | 2.5 | 2.0 | 78% | 1145 | 1057 | 69.4 |
| *R. clinocephalus* | 1.4 | 1.1 | 79% | 1120 | 979 | 38.4 |
| *R. nepalensis* | 0.3 | 0.2 | 81% | 620 | 89 | 7.6 |
| *R. nivalis* | 0.8 | 0.7 | 80% | 1097 | 910 | 23.3 |
| *R. niveus* | 1.0 | 0.6 | 58% | 1090 | 909 | 21.1 |
| *R. occidentalis* | 2.1 | 1.2 | 59% | 1135 | 1075 | 43.6 |
| *R. odoratus* | 2.2 | 1.2 | 54% | 1135 | 1034 | 41.3 |
| *R. palmatus* | 0.3 | 0.3 | 77% | 701 | 260 | 9.1 |
| *R. parviflorus* | 1.3 | 0.9 | 73% | 1130 | 1011 | 33.2 |
| *R. parvifolius* | 1.7 | 1.2 | 69% | 1129 | 1053 | 41.7 |
| *R. parvus* | 3.0 | 1.4 | 45% | 1137 | 1037 | 48.5 |
| *R. pectinarioides* | 2.5 | 2.1 | 81% | 1142 | 1086 | 72.0 |
| *R. pectinellus** | 0.0 | 0.0 | 75% | 0 | - | 0.1 |
| *R. pedatus* | 3.8 | 3.0 | 78% | 1143 | 1104 | 105.3 |
| *R. pensilvanicus* | 1.5 | 1.2 | 82% | 1127 | 1053 | 41.7 |
| *R. pentagonus* | 1.8 | 1.3 | 76% | 1130 | 984 | 46.9 |
| *R. phoenicolasius* | 2.9 | 2.3 | 82% | 1155 | 1110 | 82.4 |
| *R. pubescens* | 1.1 | 0.7 | 67% | 1110 | 956 | 25.6 |
| *R. pungens* | 0.8 | 0.5 | 66% | 1099 | 874 | 18.5 |
| *R. repens* | 1.0 | 0.7 | 74% | 1111 | 938 | 26.3 |
| *R. robustus* | 2.6 | 2.1 | 82% | 1144 | 1091 | 74.8 |
| *R. roseus* | 3.3 | 2.4 | 72% | 1146 | 1090 | 84.1 |
| *R. rosifolius* | 4.5 | 3.3 | 75% | 1145 | 1081 | 117.0 |
| *R. sachalinensis* | 4.3 | 3.2 | 75% | 1151 | 1114 | 111.8 |
| *R. saxatilis* | 4.2 | 2.6 | 62% | 1152 | 1105 | 91.9 |
| *R. sengorensis* | 5.3 | 2.2 | 41% | 1142 | 1073 | 75.5 |
| *R. setosus* | 4.7 | 2.3 | 48% | 1137 | 1090 | 79.1 |
| *R. spectabilis* | 3.6 | 2.0 | 54% | 1145 | 1098 | 69.5 |
| *R. strigosus* | 7.7 | 3.4 | 45% | 1155 | 1127 | 120.7 |
| *R. tephrodes* | 1.1 | 0.8 | 75% | 1111 | 913 | 28.3 |
| *R. thomsonii* | 0.9 | 0.7 | 70% | 1082 | 789 | 22.9 |
| *R. treutleri* | 0.9 | 0.7 | 72% | 1092 | 848 | 23.9 |
| *R. tricolor* | 0.7 | 0.4 | 58% | 1023 | 581 | 14.9 |
| *R. trifidus* | 1.5 | 0.6 | 44% | 1098 | 929 | 22.2 |
| *R. trilobus* | 2.6 | 1.7 | 68% | 1143 | 1080 | 61.5 |
| *R. trivialis* | 0.6 | 0.3 | 58% | 992 | 526 | 12.0 |
| *R. ulmifolius* | 3.0 | 2.0 | 66% | 1144 | 1095 | 69.7 |
| *R. ursinus* x *R. armeniacus 1* | 4.2 | 3.2 | 75% | 1149 | 1116 | 111.0 |
| *R. ursinus2* | 1.0 | 0.8 | 78% | 1122 | 908 | 27.1 |
| *R. ursinus3* | 5.8 | 3.5 | 61% | 1157 | 1125 | 123.8 |
| *R. ursinus4* | 2.0 | 1.7 | 85% | 1144 | 1070 | 59.4 |
| *R. ursinus5* | 2.0 | 1.5 | 76% | 1142 | 1055 | 54.1 |
| *R. ursinus* x *R. armeniacus 6* | 1.2 | 0.8 | 63% | 1100 | 871 | 27.3 |
| *R. urticifolius* | 1.4 | 0.7 | 49% | 1108 | 948 | 24.4 |
| *W. fragarioides* | 1.1 | 0.3 | 28% | 856 | 266 | 11.4 |

Additional file 2: Table S2. Alignment statistics for analyses for exon and supercontigs (S, exon + noncoding sequences) for diploid and all taxa *Rubus* datasets. Samples marked with an asterisk did not sequence well and were not included in phylogenetic analyses. E = exon sequence alignments. Ungapped lengths are reported.

| **Sample** | **% Cov. strawberry/apple/peach targets** | **% Cov. *R. occ.* targets** | **Length of S seq. alignment in All Taxa (Mbp)** | **% Non-gap bases in S seq. alignment for all taxa** | **Length of S seq. alignment in diploid taxa (Mbp)** | **% Non-gap bases in S seq. alignment for diploid taxa** | **Length of E seq. alignment in all taxa (Mbp)** | **% Non-gap bases in E seq. alignment for all taxa** | **Length of E seq. alignment in diploid taxa (Mbp)** | **% Non-gap bases in E seq. alignment for diploid taxa** |
| --- | --- | --- | --- | --- | --- | --- | --- | --- | --- | --- |
| **Average** | **86%** | **101%** | **2.7** | **27%** | **3.9** | **36%** | **1.5** | **58%** | **1.6** | **61%** |
| Boysen | 91% | 108% | 1.46 | 15% | - | - | 0.81 | 32% | - | - |
| Logan | 97% | 113% | 1.61 | 16% | - | - | 1.11 | 44% | - | - |
| Marion | 92% | 109% | 1.83 | 18% | - | - | 1.12 | 44% | - | - |
| *R. acanthophyllus* | 95% | 112% | 1.29 | 13% | - | - | 1.26 | 50% | - | - |
| *R. alexeterius* | 88% | 105% | 2.78 | 28% | 2.89 | 29% | 1.54 | 61% | 1.45 | 57% |
| *R. allegheniensis* | 18% | 27% | 0.01 | 0% | 0.03 | 0% | 0.05 | 2% | 0.04 | 1% |
| *R. amphidasys* | 96% | 111% | 1.18 | 12% | - | - | 1.38 | 54% | - | - |
| *R. arcticus* | 93% | 107% | 4.58 | 45% | 4.64 | 46% | 2.05 | 81% | 1.86 | 73% |
| *R. argutus* | 93% | 106% | 4.14 | 41% | 4.26 | 42% | 2.03 | 80% | 1.89 | 75% |
| *R. armeniacus* | 79% | 97% | 3.87 | 38% | - | - | 1.72 | 68% | - | - |
| *R. assamensis* | 98% | 112% | 0.86 | 8% | - | - | 1.20 | 47% | - | - |
| *R. australis* | 98% | 112% | 1.10 | 11% | - | - | 1.29 | 51% | - | - |
| *R. bifrons* | 90% | 103% | 3.81 | 38% | - | - | 1.88 | 74% | - | - |
| *R. caesius* | 97% | 112% | 0.93 | 9% | - | - | 0.68 | 27% | - | - |
| *R. calophyllus* | 90% | 106% | 1.17 | 12% | - | - | 1.01 | 40% | - | - |
| *R. calycinus* | 86% | 104% | 1.04 | 10% | - | - | 0.86 | 34% | - | - |
| *R. canadensis* | 95% | 107% | 3.93 | 39% | 4.11 | 41% | 1.93 | 76% | 1.83 | 72% |
| *R. caucasicus* | 85% | 99% | 3.75 | 37% | - | - | 1.74 | 69% | - | - |
| *R. chamaemorus* | 66% | 75% | 0.80 | 8% | - | - | 0.88 | 35% | - | - |
| *R. coreanus* | 95% | 110% | 4.78 | 47% | 4.85 | 48% | 1.95 | 77% | 1.76 | 69% |
| *R. coriifolius* | 78% | 94% | 3.50 | 35% | 3.57 | 35% | 1.88 | 74% | 1.64 | 64% |
| *R. crataegifolius* | 94% | 107% | 3.49 | 35% | 3.53 | 35% | 1.92 | 76% | 1.74 | 69% |
| *R. cuneifolius* | 92% | 104% | 4.53 | 45% | 4.60 | 45% | 2.09 | 83% | 1.93 | 76% |
| *R. deliciosus* | 93% | 105% | 5.13 | 51% | 5.18 | 51% | 2.11 | 83% | 1.88 | 74% |
| *R. ellipticus* | 93% | 105% | 3.16 | 31% | 3.52 | 35% | 1.43 | 57% | 1.41 | 56% |
| *R. eriocarpus* | 93% | 108% | 5.59 | 55% | 5.65 | 56% | 2.13 | 84% | 2.01 | 79% |
| *R. flagellaris* | 97% | 114% | 1.17 | 12% | - | - | 1.11 | 44% | - | - |
| *R. fockeanus* | 97% | 112% | 1.46 | 14% | - | - | 1.00 | 39% | - | - |
| *R. geoides* | 97% | 113% | 0.94 | 9% | - | - | 1.18 | 46% | - | - |
| *R. glabratus* | 96% | 113% | 1.32 | 13% | - | - | 1.34 | 53% | - | - |
| *R. glaucus* | 81% | 105% | 1.91 | 19% | - | - | 1.09 | 43% | - | - |
| *R. gunnianus* | 94% | 109% | 1.66 | 16% | - | - | 1.45 | 57% | - | - |
| *R. hawaiensis* | 86% | 102% | 4.99 | 49% | 5.06 | 50% | 2.15 | 85% | 1.90 | 75% |
| *R. hispidus** | - | - | - | - | - | - | - | - | - | - |
| *R. humulifolius* | 86% | 98% | 2.32 | 23% | - | - | 1.24 | 49% | - | - |
| *R. ichangensis* | 98% | 114% | 0.97 | 10% | - | - | 1.17 | 46% | - | - |
| *R. idaeus* | 93% | 109% | 4.61 | 46% | 4.73 | 47% | 1.93 | 76% | 1.79 | 70% |
| *R. illecebrosus* | 95% | 106% | 4.47 | 44% | 4.50 | 44% | 2.08 | 82% | 1.87 | 74% |
| *R. innominatus* | 91% | 105% | 5.11 | 51% | 5.22 | 52% | 1.93 | 76% | 1.72 | 68% |
| *R. irenaeus* | 97% | 113% | 1.37 | 14% | - | - | 1.55 | 61% | - | - |
| *R. laciniatus* | 89% | 103% | 3.64 | 36% | - | - | 1.73 | 68% | - | - |
| *R. laegardii* | 86% | 101% | 1.79 | 18% | - | - | 1.26 | 50% | - | - |
| *R. lambertianus* | 93% | 110% | 1.17 | 12% | - | - | 1.19 | 47% | - | - |
| *R. lasiococcus* | 78% | 94% | 3.23 | 32% | 3.28 | 32% | 1.76 | 70% | 1.53 | 60% |
| *R. lasiostylus* | 95% | 108% | 5.40 | 54% | 5.41 | 53% | 1.87 | 74% | 1.69 | 67% |
| *R. leucodermis* | 93% | 110% | 5.87 | 58% | 5.92 | 58% | 2.11 | 83% | 1.99 | 78% |
| *R. lineatus* | 90% | 108% | 1.36 | 14% | - | - | 1.01 | 40% | - | - |
| *R. loxensis* | 83% | 98% | 2.24 | 22% | - | - | 1.32 | 52% | - | - |
| *R. macraei* | 92% | 113% | 0.96 | 10% | - | - | 0.94 | 37% | - | - |
| *R. moorei* | 91% | 108% | 0.79 | 8% | - | - | 1.28 | 51% | - | - |
| *R. clinocephaluss* | 82% | 103% | 1.90 | 19% | - | - | 1.33 | 52% | - | - |
| *R. nepalensis* | 20% | 22% | 0.11 | 1% | - | - | 0.12 | 5% | - | - |
| *R. nivalis* | 79% | 95% | 2.89 | 29% | 3.12 | 31% | 1.38 | 54% | 1.60 | 63% |
| *R. niveus* | 75% | 96% | 3.42 | 34% | 3.44 | 34% | 1.84 | 72% | 1.57 | 62% |
| *R. occidentalis* | 87% | 107% | 5.67 | 56% | 5.75 | 57% | 2.14 | 85% | 1.97 | 77% |
| *R. odoratus* | 90% | 103% | 4.84 | 48% | 4.86 | 48% | 2.11 | 83% | 1.85 | 73% |
| *R. palmatus* | 29% | 42% | 0.77 | 8% | 0.78 | 8% | 0.56 | 22% | 0.45 | 18% |
| *R. parviflorus* | 87% | 103% | 4.21 | 42% | 4.24 | 42% | 2.02 | 80% | 1.80 | 71% |
| *R. parvifolius* | 87% | 104% | 4.49 | 44% | 4.59 | 45% | 2.02 | 79% | 1.82 | 72% |
| *R. parvus* | 90% | 106% | 1.48 | 15% | - | - | 1.33 | 52% | - | - |
| *R. pectinarioides* | 94% | 110% | 1.70 | 17% | - | - | 1.25 | 49% | - | - |
| *R. pectinellus** | - | - | - | - | - | - | - | - | - | - |
| *R. pedatus* | 95% | 108% | 4.20 | 42% | 4.25 | 42% | 1.85 | 73% | 1.68 | 66% |
| *R. pensilvanicus* | 89% | 104% | 3.54 | 35% | - | - | 1.93 | 76% | - | - |
| *R. pentagonus* | 86% | 101% | 2.14 | 21% | - | - | 1.01 | 40% | - | - |
| *R. phoenicolasius* | 93% | 110% | 4.44 | 44% | 4.52 | 45% | 1.96 | 77% | 1.73 | 68% |
| *R. pubescens* | 80% | 98% | 4.07 | 40% | 4.15 | 41% | 1.95 | 77% | 1.74 | 68% |
| *R. pungens* | 76% | 93% | 2.03 | 20% | 2.30 | 23% | 1.26 | 50% | 1.29 | 51% |
| *R. repens* | 84% | 95% | 2.23 | 22% | 2.27 | 22% | 1.42 | 56% | 1.33 | 52% |
| *R. robustus* | 94% | 106% | 4.29 | 43% | 4.37 | 43% | 2.02 | 80% | 1.87 | 74% |
| *R. roseus* | 94% | 111% | 1.39 | 14% | - | - | 1.32 | 52% | - | - |
| *R. rosifolius* | 94% | 105% | 4.53 | 45% | 4.57 | 45% | 2.07 | 81% | 1.87 | 73% |
| *R. sachalinensis* | 94% | 109% | 5.04 | 50% | - | - | 1.98 | 78% | - | - |
| *R. saxatilis* | 95% | 111% | 1.82 | 18% | - | - | 1.01 | 40% | - | - |
| *R. sengorensis* | 93% | 110% | 1.17 | 12% | - | - | 1.13 | 44% | - | - |
| *R. setosus* | 93% | 107% | 4.09 | 41% | 4.36 | 43% | 1.73 | 68% | 1.76 | 69% |
| *R. spectabilis* | 93% | 109% | 5.24 | 52% | 5.31 | 52% | 2.12 | 83% | 1.94 | 76% |
| *R. strigosus* | 96% | 111% | 5.04 | 50% | 5.14 | 51% | 2.00 | 79% | 1.81 | 71% |
| *R. tephrodes* | 78% | 96% | 1.39 | 14% | - | - | 1.25 | 49% | - | - |
| *R. thomsonii* | 71% | 86% | 2.07 | 20% | - | - | 1.02 | 40% | - | - |
| *R. treutleri* | 75% | 90% | 1.78 | 18% | - | - | 1.28 | 51% | - | - |
| *R. tricolor* | 60% | 70% | 1.17 | 12% | - | - | 0.79 | 31% | - | - |
| *R. trifidus* | 82% | 94% | 2.95 | 29% | 3.05 | 30% | 1.79 | 71% | 1.57 | 62% |
| *R. trilobus* | 93% | 107% | 5.37 | 53% | 5.40 | 53% | 2.17 | 86% | 1.97 | 78% |
| *R. trivialis* | 58% | 67% | 0.43 | 4% | 0.51 | 5% | 0.52 | 20% | 0.51 | 20% |
| *R. ulmifolius* | 93% | 106% | 3.76 | 37% | 3.86 | 38% | 2.00 | 79% | 1.80 | 71% |
| *R. ursinus* x *R. armeniacus 1* | 95% | 113% | 1.33 | 13% | - | - | 1.15 | 45% | - | - |
| *R. ursinus2* | 77% | 95% | 1.56 | 15% | - | - | 0.98 | 39% | - | - |
| *R. ursinus3* | 96% | 114% | 1.63 | 16% | - | - | 1.29 | 51% | - | - |
| *R. ursinus4* | 89% | 110% | 1.94 | 19% | - | - | 1.36 | 53% | - | - |
| *R. ursinus5* | 88% | 108% | 1.97 | 20% | - | - | 1.37 | 54% | - | - |
| *R. ursinus* x *R. armeniacus 6* | 75% | 92% | 2.81 | 28% | - | - | 1.31 | 52% | - | - |
| *R. urticifolius* | 81% | 98% | 3.54 | 35% | 3.59 | 35% | 2.03 | 80% | 1.79 | 70% |
| *W. fragarioides* | 68% | 36% | 0.02 | 0% | 0.02 | 0% | 0.07 | 3% | 0.08 | 3% |

Additional file 3: Table S3. Chloroplast sequence statistics. *Rubus* samples marked with an asterisk either failed sequencing or were removed from the analysis as rogue taxa by RogueNaRok. Average read depth was calculated from mapping up to 100,000 reads against the *R. occidentalis* reference chloroplast sequence edited to include one copy of the inverted repeat region.

| **Sample** | **Avg. read depth of chloroplast sequence** | **Ungapped length of the chloroplast genome (bp)** | **% Gaps in chloroplast genome** |
| --- | --- | --- | --- |
| **Average** | **23** | **124,184** | **1%** |
| Boysen | 12.6 | 124,309 | 1% |
| Logan | 37.7 | 125,567 | 0% |
| Marion | 21.4 | 125,663 | 0% |
| *R. acanthophyllus* | 11.1 | 122,694 | 2% |
| *R. alexeterius* | 7.9 | 123,719 | 2% |
| *R. allegheniensis* | 3.1 | 115,577 | 8% |
| *R. amphidasys* | 29.2 | 125,522 | 0% |
| *R. arcticus* | 17.5 | 123,400 | 2% |
| *R. argutus* | 16 | 125,537 | 0% |
| *R. armeniacus* | 5.6 | 125,441 | 0% |
| *R. assamensis* | 20.2 | 125,711 | 0% |
| *R. australis* | 6.9 | 125,838 | 0% |
| *R. bifrons* | 7.7 | 124,415 | 1% |
| *R. caesius* | 61.9 | 125,659 | 0% |
| *R. calophyllus* | 1.4 | 111,561 | 11% |
| *R. calycinus* | 6 | 121,047 | 4% |
| *R. canadensis* | 11.8 | 125,432 | 0% |
| *R. caucasicus** | 1.9 | - | - |
| *R. chamaemorus* | 1.9 | 125,640 | 0% |
| *R. coreanus* | 11.2 | 125,007 | 1% |
| *R. coriifolius* | 4.3 | 124,525 | 1% |
| *R. crataegifolius* | 59.1 | 125,125 | 1% |
| *R. cuneifolius* | 4.5 | 125,520 | 0% |
| *R. deliciosus* | 24.9 | 125,102 | 1% |
| *R. ellipticus* | 12.3 | 123,604 | 2% |
| *R. eriocarpus* | 9.6 | 124,179 | 1% |
| *R. flagellaris* | 12.6 | 122,233 | 3% |
| *R. fockeanus* | 48.8 | 125,693 | 0% |
| *R. geoides* | 16.6 | 125,418 | 0% |
| *R. glabratus* | 74.4 | 125,383 | 0% |
| *R. glaucus* | 6.5 | 125,225 | 0% |
| *R. gunnianus* | 32.2 | 125,184 | 0% |
| *R. hawaiensis* | 5.1 | 125,115 | 1% |
| *R. hispidus** | - | - | - |
| *R. humulifolius* | 9.2 | 123,176 | 2% |
| *R. ichangensis* | 27.1 | 125,688 | 0% |
| *R. idaeus* | 58 | 124,689 | 1% |
| *R. illecebrosus* | 68.2 | 124,484 | 1% |
| *R. innominatus* | 4 | 124,262 | 1% |
| *R. irenaeus* | 12 | 125,706 | 0% |
| *R. laciniatus* | 8.9 | 125,496 | 0% |
| *R. laegardii* | 3.5 | 118,943 | 5% |
| *R. lambertianus** | 6.5 | - | - |
| *R. lasiococcus* | 4.5 | 124,906 | 1% |
| *R. lasiostylus* | 47.9 | 124,685 | 1% |
| *R. leucodermis* | 76 | 125,721 | 0% |
| *R. lineatus* | 3 | 123,760 | 2% |
| *R. loxensis* | 2.8 | 125,068 | 1% |
| *R. macraei* | 8.7 | 121,732 | 3% |
| *R. moorei* | 5 | 124,514 | 1% |
| *R. clinocephaus* | 2.6 | 125,129 | 1% |
| *R. nepalensis* | 3.4 | 124,770 | 1% |
| *R. nivalis* | 7.1 | 121,338 | 4% |
| *R. niveus* | 14.4 | 125,501 | 0% |
| *R. occidentalis* | 24.9 | 125,769 | 0% |
| *R. odoratus* | 41.8 | 125,620 | 0% |
| *R. palmatus* | 1.6 | 125,003 | 1% |
| *R. parviflorus* | 19 | 125,587 | 0% |
| *R. parvifolius* | 14.8 | 125,386 | 0% |
| *R. parvus* | 75.5 | 125,463 | 0% |
| *R. pectinarioides* | 7.1 | 125,435 | 0% |
| *R. pectinellus** | - | - | - |
| *R. pedatus* | 49 | 125,696 | 0% |
| *R. pensilvanicus* | 7.1 | 124,309 | 1% |
| *R. pentagonus* | 3.8 | 125,580 | 0% |
| *R. phoenicolasius* | 26.3 | 125,140 | 1% |
| *R. pubescens* | 8.5 | 125,487 | 0% |
| *R. pungens* | 16.5 | 125,668 | 0% |
| *R. repens* | 3.1 | 111,902 | 11% |
| *R. robustus** | 8.3 | - | - |
| *R. roseus* | 17.1 | 125,430 | 0% |
| *R. rosifolius* | 17.8 | 124,949 | 1% |
| *R. sachalinensis* | 36.9 | 124,785 | 1% |
| *R. saxatilis* | 75.9 | 125,698 | 0% |
| *R. sengorensis* | 75.7 | 125,724 | 0% |
| *R. setosus* | 74.9 | 125,560 | 0% |
| *R. spectabilis* | 75.2 | 125,426 | 0% |
| *R. strigosus* | 74.9 | 124,818 | 1% |
| *R. tephrodes* | 8.9 | 125,337 | 0% |
| *R. thomsonii* | 14.7 | 125,710 | 0% |
| *R. treutleri* | 7.2 | 124,235 | 1% |
| *R. tricolor* | 24.3 | 125,699 | 0% |
| *R. trifidus* | 48.3 | 124,841 | 1% |
| *R. trilobus* | 52 | 125,283 | 0% |
| *R. trivialis* | 11 | 125,687 | 0% |
| *R. ulmifolius* | 38 | 125,701 | 0% |
| *R. ursinus* x  *R. armeniacus 1* | 34.5 | 125,664 | 0% |
| *R. ursinus2* | 1.3 | 106,708 | 15% |
| *R. ursinus3* | 74.8 | 125,597 | 0% |
| *R. ursinus4* | 11.7 | 125,035 | 1% |
| *R. ursinus5* | 8.5 | 125,496 | 0% |
| *R. ursinus* x  *R. armeniacus* *6* | 10.9 | 125,604 | 0% |
| *R. urticifolius* | 38.7 | 125,619 | 0% |
| *W. fragarioides* | 15.8 | 116,194 | 8% |

Additional file 4: Table S4. Putative *Rubus* hybrid groups and species. Hybrid origins supported by discordance in nuclear phylogenies and networks. Maternal progenitors inferred from the chloroplast phylogeny.

| **Putative Hybrids** | **Putative parental groups or taxa** |
| --- | --- |
| Group 5 | Putative parents from Group 7 and Groups 3 and 4 (Possibly *Idaeobatus* and *Cylactis*) |
| Group 6 | Subg. *Rubus* $\times$ *Idaeobatus* hybrids |
| *R. macraei* | Subg. *Rubus* $\times$ *Idaeobatus* hybrid |
| *R. glaucus* | Subg. *Rubus* $\times$ *Idaeobatus* hybrid |
| *R. caesius* | Subg. *Rubus* $\times$ *Idaeobatus* hybrid |
| *R. ursinus* | Subg. *Rubus* $\times$ *Idaeobatus* hybrid |
| *R. ursinus* 1 and 6 | *R. ursinus* $\times$ *armeniacus* hybrid |
| *R. humulifolius* | *Idaeobatus* $\times$ *Cylactis* hybrid |
| *R. saxatilis* | Unknown; Maternal parent likely from *Idaeobatus* and may be a black raspberry |
| *R. pentagonus, R. sengorensis, R. thomsonii* | *Idaeobatus* hybrids or closely related to *Malachobatus* and *Dalibardastrum* progenitors |
| 'Logan' | 'Aughinbaugh' (*R. ursinus*) $\times$ 'Red Antwerp' (*R. idaeus*) |
| 'Boysen' | 'Logan' $\times$ 'Austin Mayes' (*R. baileyanus* $\times$ *R. argutus*). *Idaeobatus* is grandparent via 'Logan' |
| 'Marion' | 'Youngberry' $\times$ 'Olallie'. Derived from *R. ursinus* related cultivars. *Rubus idaeus* multiple generations back in pedigree. |

Additional file Table S5. Number of studied accessions classified in each subgenus. "Other hybrids" include two *R. ursinus* × *armeniacus* hybrids.

| **Subgenus** | **Accessions** |
| --- | --- |
| *Anoplobatus* | 4 |
| *Chamaebatus* | 5 |
| *Chamaemorus* | 1 |
| *Comaropsis* | 1 |
| *Cylactis* | 7 |
| *Dalibarda* | 1 |
| *Dalibardastrum* | 4 |
| *Diemenicus* | 1 |
| *Idaeobatus* | 24 |
| *Malachobatus* | 8 |
| *Micranthobatus* | 3 |
| *Orobatus* | 5 |
| *Rubus* | 24 |
| Horticultural Hybrid | 3 |
| Other Hybrid | 2 |
| Outgroup | 1 |
| **Total** | **94** |
